# The serine-threonine kinase TAO3 promotes cancer invasion and tumor growth by facilitating trafficking of endosomes containing the invadopodia scaffold TKS5α

**DOI:** 10.1101/2020.02.27.968305

**Authors:** Shinji Iizuka, Manuela Quintavalle, Jose Ceja Navarro, Kyle P. Gribbin, Robert J. Ardecky, Matthew Abelman, Chen-Ting Ma, Eduard Sergienko, Fu-Yue Zeng, Ian Pass, George Thomas, Shannon McWeeney, Christian A. Hassig, Anthony B Pinkerton, Sara A Courtneidge

## Abstract

Invadopodia are actin-based proteolytic membrane protrusions required for invasive behavior and tumor growth. We used our high-content screening assay to identify kinases impacting invadopodia formation. Among the top hits we selected TAO3, a STE20-like kinase of the GCK subfamily, for further analysis. TAO3 was over-expressed in many human cancers, and regulated invadopodia formation in melanoma, breast and bladder cancers. Furthermore, TAO3 catalytic activity facilitated melanoma growth in 3-dimensional matrices and *in vivo*. We developed potent catalytic inhibitors of TAO3 that inhibited invadopodia formation and function, and tumor cell extravasation and growth. Using these inhibitors, we determined that TAO3 activity was required for endosomal trafficking of TKS5α, an obligate invadopodia scaffold protein. A phosphoproteomics screen for TAO3 substrates revealed the dynein subunit protein LIC2 as a relevant substrate. Knockdown of LIC2 or expression of a phosphomimetic form promoted invadopodia formation. Thus, TAO3 is a new therapeutic target with a distinct mechanism of action.

**SIGNIFICANCE:** Targeting tumor invasive behavior represents an understudied opportunity. We used an unbiased screening approach to identify kinases required for invadopodia formation and function. We validated TAO3, both genetically and with a novel inhibitor, and determined TAO3 function. Our data support clinical development of this class of target.

## INTRODUCTION

Much progress has been made in recent years in the development of novel cancer therapeutics. Among small molecules, kinase inhibitors have met with particular clinical success, with approvals of agents targeting mutated “driver” kinases, as well as non-mutated but essential enzymes [1]. Despite these successes, both intrinsic and acquired resistance limit long-term efficacy. This has led to the development of inhibitors specifically targeting resistance mechanisms, as well as to combination therapies. In the case of mutant B-RAF driven melanoma, combination therapy with the B-RAF inhibitor, dabrafenib, and the MEK inhibitor, trametinib, was recently approved [2–4]. This regimen results in improved survival, although resistance emerges in most patients after approximately 1 year. There are currently no kinase inhibitor strategies for those melanomas with wild-type B-RAF. Exciting progress has also been made in immunotherapy approaches to treat cancer, with noted successes for antibodies targeting PD-1 and CTLA-4 in all subtypes of melanoma [5]. Nevertheless, resistance to these agents also limits long-term efficacy. Thus, it is important to continue to identify new therapeutic approaches for eventual use in combination with existing agents. We reasoned that identifying targets in pathways regulating aspects of the cancer phenotype distinct from cell cycle progression and immune evasion might be such an opportunity.

We decided to focus on the invasive behavior of cancer cells. Invasion is required for cancer cells to move into and out of the bloodstream (intra- and extravasation), and therefore underlies the metastasis that is responsible for most cancer deaths. These steps will have already occurred prior to diagnosis in patients for whom removal of the primary tumor is not curative, and therefore therapeutic intervention for intra- and extravasation may not be beneficial. However, there are many examples where invasive behavior has been linked to tumor growth in both primary and metastatic sites, as well as chemoresistance [6–10]; intervention in these processes would be expected to have therapeutic benefit.

One prominent mechanism by which tumor cells exhibit invasive behavior is by the formation of membrane protrusions known as invadopodia [11–13]. These complex structures coordinate the actin cytoskeleton with pericellular proteolysis, and metallo-, cysteine and serine protease activities have all been described at invadopodia [14]. It has long been known that the presence of invadopodia correlates with invasive behavior in a number of cancer cell types *in vitro*. More recently, invadopodia were observed in human cancer specimens *ex vivo* [15], and were shown to be required for tumor cell intra- and extravasation in model systems [16, 17]. Tumor expression analysis has revealed a correlation between high expression of the obligate invadopodia scaffold protein TKS5α and worse outcomes in glial-derived tumors, non-small cell lung cancer and breast cancer [6, 18, 19], as well as with increasing stage in prostate cancer and melanoma [7, 20]. Most importantly, the use of 3-dimensional (3D) culture systems and xenograft assays has revealed that invadopodia also promote the growth of tumor cells [6, 7, 21]. While the full mechanistic details of this phenotype remain to be established, it seems likely that the pericellular proteolytic activity associated with invadopodia both remodels the extracellular matrix (ECM) to the benefit of the tumor, as well as processes and activates growth factors and cytokines required to create a growth-promoting microenvironment. For example, the angiogenesis-eliciting vascular endothelial growth factor (VEGF) is processed by metalloproteases into pro-tumorigenic forms [22], and there is some evidence that loss of invadopodia results in reduced tumor-associated VEGF [6].

What might be good therapeutic targets for invadopodia inhibition? The TKS5α scaffold itself, as well as many other key invadopodia proteins, have no catalytic activity, and thus developing drugs against them, while not impossible, would be challenging [23]. Other key players, such as small GTPases, have to-date proven largely intractable to inhibitor strategies. And inhibitors of the matrix metalloproteases have also not met with success to date, perhaps because of redundancy among different protease classes, as well as lack of specificity and potency. Some years ago, we established a high-content screening assay to identify in an unbiased way regulators of invadopodia formation and function [24]. Using a small compound library in a proof-of-principle screen, we identified several small molecules annotated as cyclin-dependent kinase (CDK) inhibitors. We deconvoluted these hits to identify and subsequently validate CDK5 as their target, and provided the mechanism by which CDK5 regulates invadopodia formation [24]. Interestingly, at around the same time it was recognized that CDK5 is over-expressed and indeed frequently amplified in pancreatic cancer, which elaborate invadopodia [25]. Knockdown of CDK5 had no effect on growth of pancreatic cells on tissue culture plastic, but did markedly inhibit the growth of the cells in 3D matrices as well as *in vivo*, in keeping with a role for CDK5 as an invadopodia regulator [26]. Together these studies support the conclusion that invadopodia inhibition is a viable strategy to reduce tumor growth, and suggest that kinases might be a valuable class of invadopodia targets. Despite the value of kinase inhibitors for cancer therapy generally, and the large target class (there are more than 500 members of the kinome [27]), it is quite remarkable that most research, as well as most target identification and validation [28], and inhibitor development [29], have focused on just a few enzymes. We therefore set out to screen the entire kinome to identify invadopodia regulators.

## RESULTS

### Screening for kinases regulating invadopodia formation

We previously described a high-content image-based screening assay designed to identify regulators of invadopodia formation [24]. Our strategy was to use Src3T3 cells (a mouse model of fibrosarcoma that elaborates abundant invadopodia) for the initial screen. Top hits were subsequently validated in human cancer cell lines. This study led to the identification and validation of CDK5. We used the same assay here. A library of pooled siRNAs (3 for each murine kinase [30]), targeting all protein kinases as well as select lipid and metabolic kinases, was kindly provided by Dr. Pam Carroll of Merck. Src3T3 cells, in 1536 well format, were transfected with the pools, incubated for 48 hours, then stained with phalloidin and DAPI to label the F-actin in invadopodia and nuclear DNA respectively. Following automatic well focusing and image capture, each well was analyzed by eye for cell number and viability. For those wells where no cytotoxicity or apoptosis was observed, invadopodia were then evaluated. From this, 12 kinases were prioritized. One of these was the previously validated CDK5, which we did not explore further here. For the others, we next tested each siRNA separately, requiring at least 2 of the 3 to inhibit to proceed. One kinase was deemed off-target by this criterion, leaving 10 in our list (Figure 1A). Representative images for these are shown in Figure 1B and the results summarized in Table 1. Next, we performed a functional assay (gelatin degradation). Knockdown of 3 of these kinases, while inhibiting invadopodia formation, did not affect matrix degradation (Figure 1C, Table 1), and were not pursued further.

**Figure 1.**
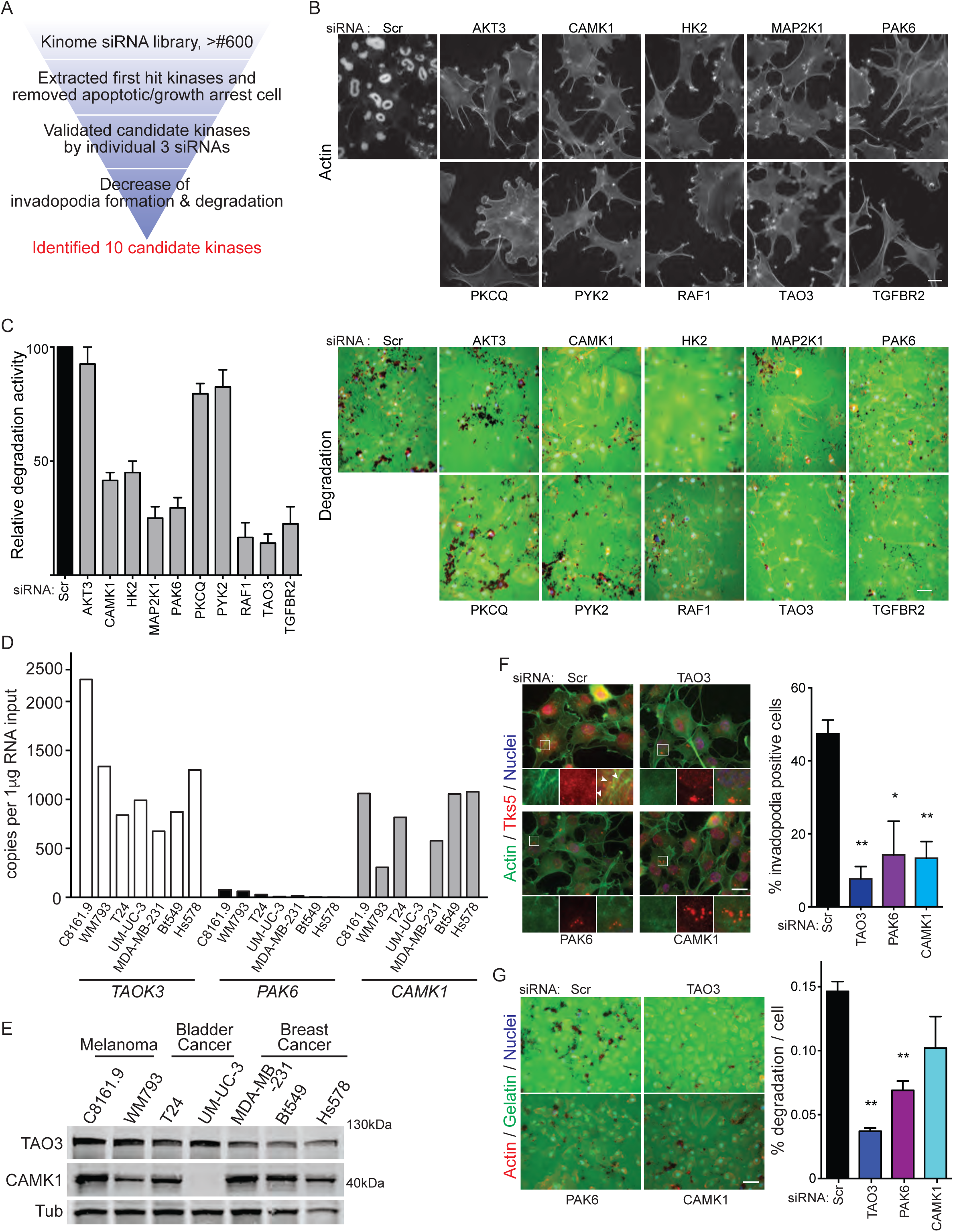
High-content screening image analysis using a kinome siRNA library reveals invadopodia regulators and validation of candidate kinases. **A** Schematic view of screening and validation steps. **B** and **C** Validation analysis of candidate kinases in Src3T3 cells. Invadopodia formation (**B**) and gelatin degradation (**C**) with siRNA-scrambled and other 10 candidate kinases. Data shown are representative images and relative degradation activity (**C**, graph). Immunofluorescence staining of invadopodia by actin (phalloidin in B and C) and gelatin (green in C). **D** and **E** Expression of top hit kinases. qPCR (**D**) and immunoblotting (**E**) analysis of TAO3, PAK6 and CAMK1 in melanoma, bladder cancer and breast cancer cell lines. Tubulin is shown as a loading control. Invadopodia formation (**F**) and gelatin degradation (**G**) analysis in C8161.9 cells with siRNA-scrambled, -TAO3, -PAK6 or –CAMK1. Representative images (left) and percentage of invadopodia positive cells or percentage of degradation per cells (right). Immunofluorescence staining of invadopodia by actin (phalloidin, green in F and red in G) and TKS5 (red in F), gelatin (green in G) and Hoechst to denote nuclei of cells. Arrowheads indicate TKS5+ invadopodia. Scale bars, 10 μm (B), 20μm (F) and 50 μm (C, G). Screening shown was run twice with quadruplicates each time. qPCR data shown are technical duplicates and were validated in 2 separated experiments. Immunoblotting data shown was validated in 2 separate experiments. Invadopodia and degradation assay shown in F and G is n=5 to 7 in each group and 2 or more biological replicates. *P*>0.05 unless other specified; *, *P*<0.05; **, *P*<0.01.

**Table 1.** Summary of screening for kinases regulating invadopodia formation and function. 10 kinases of top hit. Invadopodia formation assay was performed by pooled siRNA and individual three siRNAs. Degradation assay was performed by pooled siRNA.

| GENE | Protein Name | Invadopodia assay |  |  |  | Degradation assay |
| --- | --- | --- | --- | --- | --- | --- |
|  |  | Pool | siRNA#1 | siRNA#2 | siRNA#3 | Pool |
| CAMK1 | calcium/calmodulin-dependent protein kinase type 1 | + | + | + | + | + |
| MAP2K1 | mitogen-activated protein kinase kinase 1 | + | X | + | + | + |
| PAK6 | serine/threonine-protein kinase PAK 6 | + | + | + | + | + |
| RAF1 | RAF proto-oncogene serine/threonine-protein kinase | + | + | + | + | + |
| TAOK3 | serine/threonine-protein kinase TAO3 | + | + | + | + | + |
| TGFBR2 | TGF-beta receptor type 2 | + | + | + | X | + |
| AKT3 | RAC-gamma serine/threonine-protein kinase | + | X | + | + | X |
| PRKCQ | protein kinase C theta | + | + | + | + | X |
| PTK2 | protein-tyrosine kinase 2-beta | + | + | + | + | X |
| HK2 | hexokinase-2 | + | + | + | + | + |

Our top hits were thus narrowed to CAMK1, HK2 (a metabolic enzyme), MAP2K1, PAK6, RAF1, TAO3 and TGFβR2. Of these, we chose to focus on those kinases we considered understudied, that is with no or few reports to date on involvement in cancer progression and/or invasion. This narrowed our list down to CAMK1, PAK6 and TAO3, all serine-threonine kinases.

### Expression analysis of top hits

The next step was to determine the expression of these 3 kinases in cancer tissue and cell lines. We first performed Cancer Outlier Profile Analysis (COPA) [31] on public gene expression data sets curated by Oncomine [32]. Genes scoring in the top 10% of COPA scores at any of three percentile cutoffs (75th, 90th, and 95th) were deemed outliers in their respective datasets. At the 10% gene rank, TAOK3 was an outlier in 28 studies in 17 cancer types, PAK6 was an outlier in 31 studies in 14 cancer types and CAMK1 was an outlier in 44 studies in 16 cancer types. Some of these tumor types (eg melanoma, breast, bladder) have been associated with invadopodia formation in several studies, whereas others (particularly leukemia and lymphoma) have not been well studied. Since it was important that we be able to assess the role of these kinases in human cancer cells *in vitro*, we chose to focus on melanoma, breast and bladder cancers (Supplementary Table S1, S2, S3).

We first determined the expression of the 3 kinases in representative melanoma, bladder and breast cancer cell lines by qPCR, and for TAO3 and CAMK1 also by immunoblotting (Figure 1D, E). All kinases were expressed in the melanoma cell lines, so we chose to pursue them further in this cancer type. We first used transient RNA interference to reduce expression of each in C8161.9 cells, and evaluated both invadopodia formation and function (Figure 1F, G). The invadopodia formation assay was also performed in another human melanoma cell line, WM793, which confirmed reduction in invadopodia number (Supplementary Figure S1A). The most robust effect in both assays was observed for TAO3 knockdown. In our screening procedure, we excluded kinases that appeared to reduce cell viability. By using siRNA to knockdown each member of the TAO family in C8161.9 cells, and evaluating the cells 3 days later (Figure S1B), we found that cell number was compromised by loss of TAO1 or TAO2, but loss of TAO3 had no effect, confirming our screening data.

Finally, we determined the expression of TAO3 at the protein level in clinical specimens of melanoma. An automated immunohistochemical staining protocol was developed then used to stain 20 samples of primary melanoma. The slides were scored (using a 0-2 scale, where 0=negative; 1=weak; 2=mod-strong) and intensity evaluated across each tumor to derive an immunoscore with a range of 0-200. Nineteen of twenty evaluable samples showed staining in some portion of the tumor, with 7 with scores of 101-150 and 8 with scores greater than 151, suggesting increased expression in melanomas. In some cases, particularly strong staining could be seen at the tumor border, and in disseminating melanoma cells (Supplementary Figure S1C). Together, these analyses led us to nominate TAO3 as our lead kinase.

The gene expression analyses we described earlier suggested that TAO3 was also highly differentially expressed in breast and bladder carcinomas, and was present in cell lines derived from these tumors (Figure 1D, 1E, Supplementary Table S1). To determine the generality of our findings, we therefore tested whether TAO3 was required for invadopodia formation and function in these cells. Both invadopodia formation and gelatin degradation were inhibited (Supplementary Figure S1D, S1E – note that T24 could not be evaluated for gelatin degradation because of matrix tears caused by pulling). In addition, we determined that the TAO3 siRNA did not affect the expression of TAO1 or TAO2 (Supplementary Figure S1F). Together, these data define TAO3 as a regulator of invadopodia formation and invasion, and further reveal its importance in multiple tumor cell types.

### Validation of TAO3 in melanoma

To further validate TAO3 in melanoma, we generated a lentivirus expressing shRNA specific for TAO3, as well as non-targetable expression constructs for wildtype and kinase-dead (via mutation of lysine 53 in the ATP binding site of the catalytic domain) TAO3. Quantitative PCR demonstrated the specificity of the knockdown and successful re-expression of the constructs (Supplementary Figure S2A). Furthermore, employing a similar knockdown and re-expression strategy, we found that the catalytic activity of TAO1 was not required to rescue the cytotoxic effects seen upon TAO1 knockdown (Supplementary Figure S2B). For the TAO3 experiments, analysis of invadopodia formation (Figure 2A) and function (Figure 2B) as well as invasion through matrigel (Figure 2C) in C8161.9 melanoma cells revealed that TAO3 knockdown has a significant inhibitory effect in each case, which could be rescued by re-expression of wildtype but not kinase-inactive TAO3. We have previously shown that reducing the expression of the obligate invadopodia scaffold protein TKS5 has an inhibitory effect on growth in 3D collagen matrices, as well as *in vivo* [6, 7]. We next determined the effect of TAO3 knockdown in these same assays. When the cells were cultured on top of native type I collagen (collagen-I), no growth differences were seen. However, cells in which TAO3 expression was reduced did not grow as well in collagen-I as control cells. Growth was rescued by re-expression of wildtype, but not kinase inactive, TAO3 (Figure 2D). We also observed similar growth differences in a tumor spheroid assay (Figure 2E), where spheroid size is directly correlated with cell number (not shown). Similar effects on invadopodia formation and 3D growth were seen in a second melanoma cell line, WM793 (Supplementary Figure S2C, S2D).

**Figure 2.**
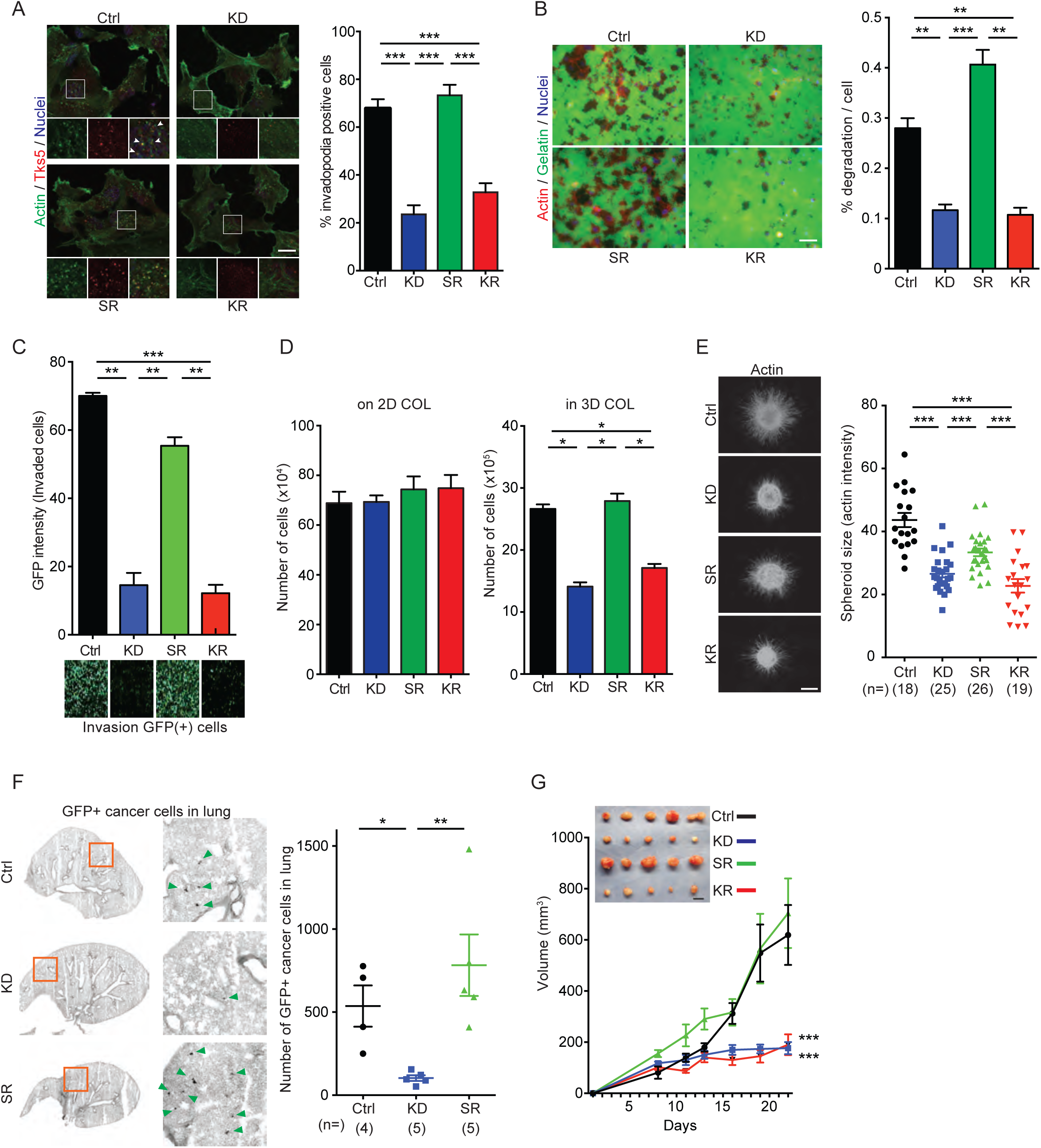
TAO3 is a regulator of invadopodia formation, invasion and growth in 3D conditions. **A** and **B**, Invadopodia formation (**A**) and gelatin degradation (**B**) analysis in C8161.9 cells with shRNA-scrambled (Scr), shRNA-TAO3 (KD), shRNA-TAO3+rescued expression of shRNA-resistant TAO3 (SR) or shRNA-TAO3+rescued expression of shRNA-resistant kinase-dead TAO3 (KR). Representative images (left) and percentage of invadopodia positive cells or percentage of degradation per cells (right). Immunofluorescence staining of invadopodia by actin (phalloidin, green) and TKS5 (red), and Hoechst to denote nuclei of cells. Arrowheads indicate TKS5+ invadopodia. **C** Boyden-chamber invasion assay with matrigel in the cells as indicated. Graph of GFP signal intensity from invaded cells (top) and representative GFP positive cells (bottom) on bottom of the membrane. **D** Growth of cells as indicated in the figure on 2D type I collagen (on 2D COL, day 8) and in 3D type I collagen (in 3D COL, day 12). **E** 3D growth/invasion in a hanging droplet spheroid assay. Representative images of spheroids in type I collagen stained by phalloidin (left) and spheroid size measured by actin intensity (right). **F** Extravasation efficiency assay in mice. Representative images of GFP+ cancer cells in lung (left) and number of GFP+ cancer cells in lung (right). Note: Blood vessels in lung tissue have high GFP background signal. **G** Tumor growth in mice injected subcutaneously. Macroscopic view of all tumors is shown (top). Scale bars, 20 μm (A), 50 μm (B) and 500 μm (E). Data shown are n=3 to 9 in each experimental group (unless other specified in figure) and were validated in 2 or more separated experiments. *P*>0.05 unless other specified; *, *P*<0.05; **, *P*<0.01; ***, *P*<0.001.

We next evaluated the effect of TAO3 knockdown *in vivo*. For this, we used 2 distinct assays. First, it has been reported that extravasation of tumor cells into the lungs requires TKS5-dependent invadopodia formation [17]. To test if this was also the case for TAO3, we introduced GFP into the control, knockdown and rescue cells to mark them, then injected them into the tail veins of immunocompromised mice. One day later, mice were sacrificed, lungs removed and sectioned, then fluorescence microscopy used to enumerate tumor cells. TAO3 knockdown markedly inhibited extravasation, which was restored by re-expression of the wildtype protein (Figure 2F). Next, the control and TAO3 knockdown cells, as well as knockdown cells re-expressing wildtype or kinase-inactive TAO3, were injected subcutaneously into immuno-compromised mice and tumor growth evaluated over time. TAO3 knockdown caused a profound inhibition of tumor growth, which was rescued by the wildtype but not kinase-inactive TAO3 (Figure 2G). These data are consistent with the growth inhibitory effects we observed in 3D cultures.

### TAO3 inhibitor identification and testing

Our rescue studies had shown that the kinase activity of TAO3 was required for its role in invadopodia formation *in vitro* and tumor progression *in vivo*. To test the therapeutic potential of TAO3 we therefore initiated a high-throughput screening campaign to identify small molecule inhibitors of the TAO3 kinase domain. Briefly, we screened a number of compound libraries including a set of 800 known kinase inhibitors, as well as 4800 compounds with similar structural features to known kinase inhibitors. We identified a number of hits, of which a series of oxindoles were the most promising, based on potency against the TAO3 kinase domain and moderate selectivity against a broad panel of kinases. A preliminary round of chemistry around the hits identified SBI-581 and SBI-029 as a proof of concept compounds (Supplementary Figure 3A). Both display good potency against TAO3 (IC_50_=42 nM and 90 nM respectively) and moderate, but largely non-overlapping selectivity against a broad panel of kinases (Supplementary Table S4). In addition, we measured the pharmacokinetics (PK) of SBI-581 in mice. While oral bioavailability was poor (%F<5), SBI-581 displayed reasonable PK after IP injection (t_1/2_=1.5 hr; AUC= 1202 hr*ng/mL; Cmax= ∼2 μM after a 10 mg/kg dose).

We first used SBI-581 in our short-term assays of invadopodia formation (Figure 3A) and gelatin degradation (Figure 3B). We observed a dose-dependent inhibition in both cases, with an EC_50_ of <50nM, a dose that had no effect on cell viability, even after several days (Supplementary Figure S3B). Similar results were obtained with SBI-029 (Supplementary Figure S3C). We also evaluated spheroid growth, and observed an inhibitor-dependent decrease after 2 days in 3D culture (Figure 3C). Then we determined the effect of SBI-581 *in vivo*, using both the extravasation and tumor growth assays described earlier. For the tumor growth assay, C8161.9 cells were implanted subcutaneously, and 10 days later, when tumors had reached approximately 80 mm^3^, daily intraperitoneal dosing at 10 mg/kg was begun. We noted a profound inhibition of tumor growth (Figure 3E) with no significant effect on body weight (Supplementary Figure S3D). For the extravasation assay, mice were injected intraperitoneally with 30 mg/kg SBI-581 (a dose chosen to achieve a projected plasma concentration of >6 μM, a concentration well in excess of the IC_50_ for TAO3), then 30 minutes later GFP-labeled melanoma cells were introduced via the tail vein. Extravasated tumor cells in the lung were quantified one day later as described earlier. SBI-581 pre-treatment significantly inhibited extravasation, compared to the DMSO control (Figure 3D). Together, these data support the validation of TAO3 as an invadopodia, invasion and growth inhibitor.

**Figure 3.**
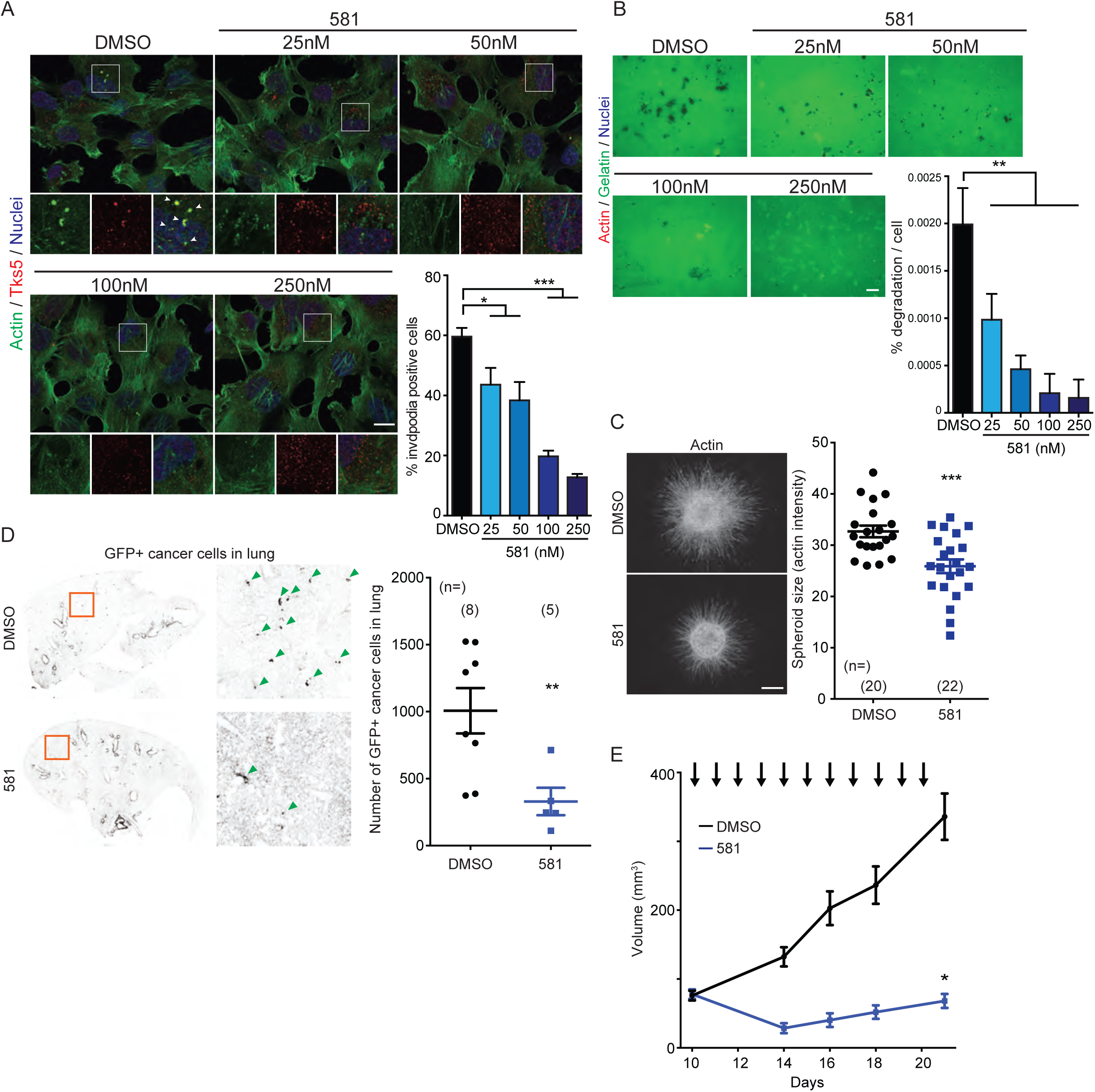
TAO3 inhibitor treatment reduces invadopodia function, 3D growth and extravasation in vitro and in vivo. **A** and **B**, Invadopodia formation (**A**) and gelatin degradation (**B**) analysis in C8161.9 cells with control DMSO and TAO3 inhibitor, SBI-581 (581). Representative images (top left) and percentage of invadopodia positive cells or percentage of degradation per cells (bottom right). Immunofluorescence staining of invadopodia by actin (phalloidin, green) and TKS5 (red), and Hoechst to denote nuclei of cells. Arrowheads indicate TKS5+ invadopodia. **C** 3D growth/invasion by hanging droplet spheroid assay. Representative images of spheroid in type I collagen stained by phalloidin (left) and spheroid size measured by actin intensity (right). **D** Extravasation efficiency assay in mice. Representative images of GFP+ cancer cells in lung (left) and number of GFP+ cancer cells in lung (right). Note: Blood vessels in lung tissue have high GFP background signal. **E** Tumor growth in mice with TAO3 inhibitor. C8161.9 cells were injected subcutaneously, then DMSO or SBI-581 (581, 10 mg/kg) was intraperitoneally injected daily (arrows). Scale bars, 20 μm (A), 50 μm (B) and 500 μm (C). Data shown are n=6 to 16 in each experimental group (unless otherwise specified in figure) and were validated in 2 or more separate experiments. *P*>0.05 unless other specified; *, *P*<0.05; **, *P*<0.01; ***, *P*<0.001.

### TAO3 and endosome trafficking

It is very important to determine how TAO3 regulates invadopodia formation, both to support its future clinical development, and to identify biomarkers to track its activity. As a first step, we determined its subcellular localization. In Src3T3 cells (which have rosettes of invadopodia), we noticed that much of the signal was not at invadopodia, but rather in a perinuclear location reminiscent of endosomes (Figure 4A). Indeed, GFP-tagged TAO3 expressed in Src3T3 cells co-localized with RAB11, a marker of recycling endosomes (Supplementary Figure S4A) [33, 34]. Endosomes are transported along a microtubule network [35], and co-staining experiments revealed a network of microtubules feeding into the invadopodia (Figure 4A) consistent with other studies [36]; TAO3 was present on these microtubules. We used high-resolution microscopy to confirm that TAO3 and RAB11 co-localized on cytoplasmic microtubules (Figure 4B). There are no reports in the literature on the involvement of RAB11 in invadopodia formation and function. Therefore we next used RNA interference to reduce its expression, which resulted in significant reduction in invadopodia number and gelatin degradation (Figure 4C).

**Figure 4.**
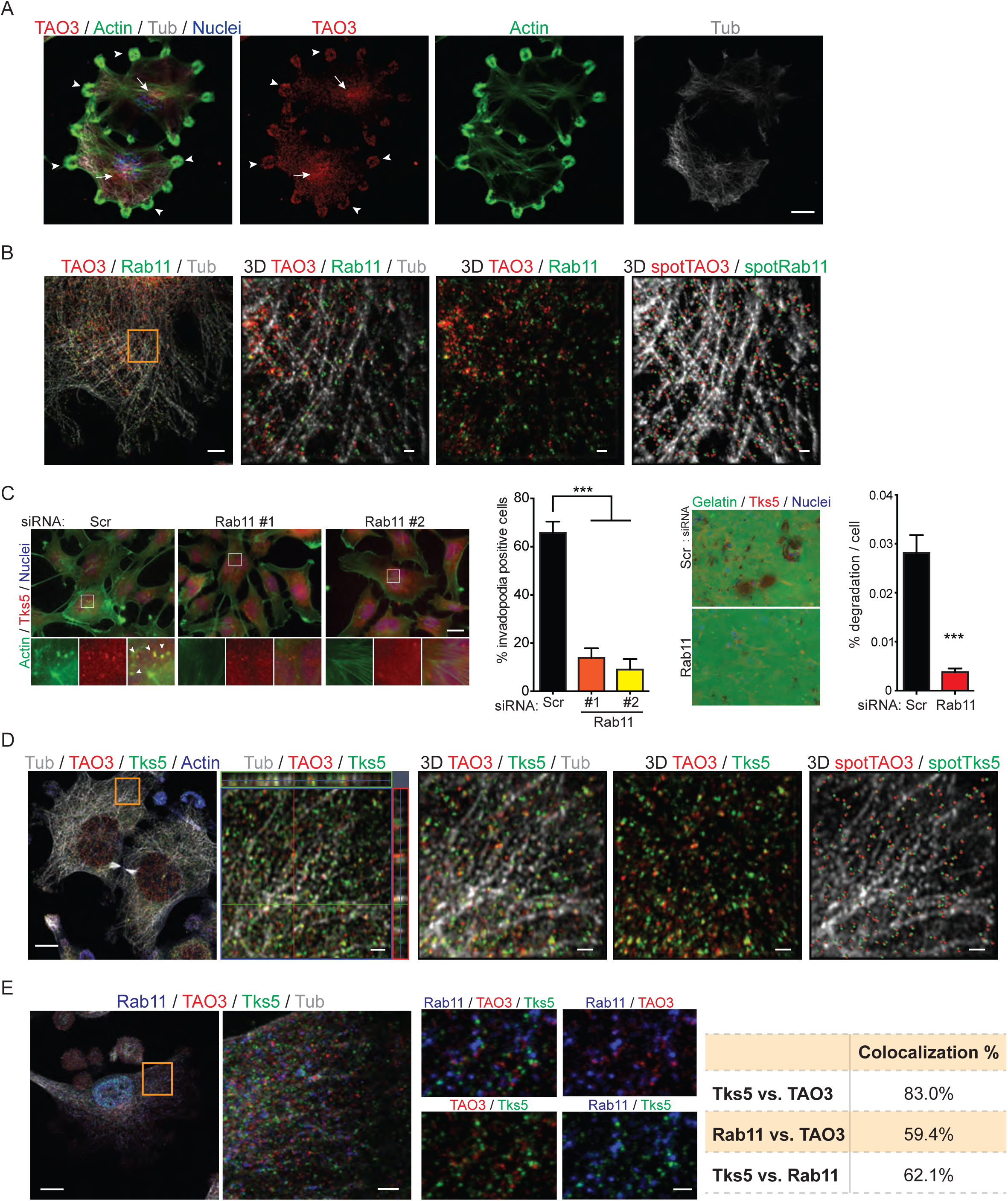
TAO3 localizes at RAB11+ endosomal vesicles with TKS5. **A** Distribution of TAO3. Staining of TAO3 (red), actin (green), tubulin (gray) and nuclei (blue in merged image only) in Src3T3 cells. Images were processed by maximum intensity projection. Arrowheads indicate invadopodial positioning of TAO3. Arrows indicate endosomal positioning of TAO3. **B** Colocalization of TAO3 and RAB11. Staining of TAO3 (red), RAB11 (green) and tubulin (gray) in Src3T3 cells. Images were processed by maximum intensity projection (left) and 3D reconstruction using Imaris software (3D, magnified area from orange square in left image). Colocalized TAO3 and RAB11 were spotted and shown with tubulin (3D spot, right). **C** Invadopodia formation (left) and gelatin degradation (right) analysis in C8161.9 cells with siRNA-scrambled and –RAB11. Two individual siRNA-RAB11 (#1 and #2) were used for invadopodia assay. Pool-siRNA-RAB11 (four siRNAs) was used for degradation assay. Representative images (left) and percentage of invadopodia positive cells or percentage of degradation per cells (right). Immunofluorescence staining of invadopodia by actin (phalloidin, green) and TKS5 (red), and Hoechst to denote nuclei of cells. Arrowheads indicate TKS5+ invadopodia. **D** Colocalization of TAO3 and TKS5*α*. Staining of tubulin (gray), TAO3 (red), TKS5*α* (green) and actin (blue) in Src3T3 cells. Images were processed by maximum intensity projection (left). Magnified area (orange square in left) was shown as orthogonal view (second left) and 3D reconstruction by Imaris software (3D). Colocalized TAO3 and TKS5*α* were spotted and shown with tubulin (3D spot, right). **E** Colocalization of RAB11, TAO3 and TKS5*α*. Staining of tubulin (gray), RAB11 (blue), TAO3 (red) and TKS5*α* (green) in Src3T3 cells. Images were processed by maximum intensity projection (left). Magnified area (orange square in left) shown (second left, single plane z-stack image) and higher magnification images are shown in small four panels with different combination of channels. Colocalization between each RAB11, TAO3 and TKS5*α* was analyzed by Imaris software and shown in table (right). Scale bars, 10 μm (A, B left, D left and E left), 1 μm (B and D right three), 2 μm (E second left) and 20 μm (C). Data shown in panel C is n=6 to 7 in each experimental group and were validated in two separate experiments. *P*>0.05 unless other specified; ***, *P*<0.001.

Our next task was to determine how RAB11 and TAO3 might impact invadopodia formation. We thought first of the invadopodia scaffold protein TKS5α [23, 37], since it has an amino-terminal PX domain with specificity for PI3P and PI3,4P_2_ [38], and PI3P in particular is highly enriched in endosomes [39, 40]. Indeed, we have recently found, using high-resolution microscopy, that TKS5α is present both at invadopodia and on microtubules (Iizuka et al, manuscript in submission, an example shown in Supplementary Figure S4B). Here we show that on microtubules, TKS5α co-localized with both TAO3 and RAB11 (Figure 4D, 4E). Furthermore, inhibition of invadopodia formation by treatment of the cells either with the Src family kinase inhibitor SU11333, or the TAO3 inhibitor SBI-581, promoted TKS5α accumulation at RAB11-positive vesicles (Supplementary Figure S4C).

We next determined the effect of the TAO3 inhibitor on endosomal trafficking of TKS5α. First, cells were engineered to express mCherry-labeled TKS5α and YFP-labeled tubulin, and trafficking was confirmed by time-lapse confocal microscopy (Figure 5A, Supplementary Movie 1). Then, either vehicle control (DMSO) or 100 nM SBI-581 was added to the cells and both time trajectory and vesicle displacement length were evaluated (Figure 5B, C, Supplementary Movie 2). The TAO3 inhibitor had a rapid and profound effect on both properties, reducing the population of motile TKS5α-positive vesicles from 24% to 6% of total. Similar effects were seen with SBI-029 (Supplementary Figure S5A). To determine if SBI-581 affected all RAB11-positive vesicles, the experiment was repeated using cells expressing DsRed-tagged RAB11. In this case, approximately the same fraction of all vesicles was motile (23%), and more than half of these retained motile characteristics after inhibitor treatment (Figure 5C, Supplementary Movie 3).This effect appeared specific to TAO3 inhibition, since inhibiting Src family kinases with SU11333 was not inhibitory (Supplementary Figure S5B, Supplementary Movie 4). Finally, SBI-581 and SBI-029 share very few targets in common (Supplementary Table S4), but we did note that one such co-inhibited kinase was ROCK2, which has noted roles in control of the actin cytoskeleton [41], and is possibly involved in invadopodia activity in response to matrix [42]. To determine if ROCK2 played any role in invadopodia formation and endocytic trafficking of TKS5α, we used the ROCK inhibitor Fasudil. ROCK inhibition affected neither invadopodia formation (Supplementary Figure S5C) nor the endocytic trafficking of TKS5α-positive vesicles (Supplementary Figure S5D).

**Figure 5.**
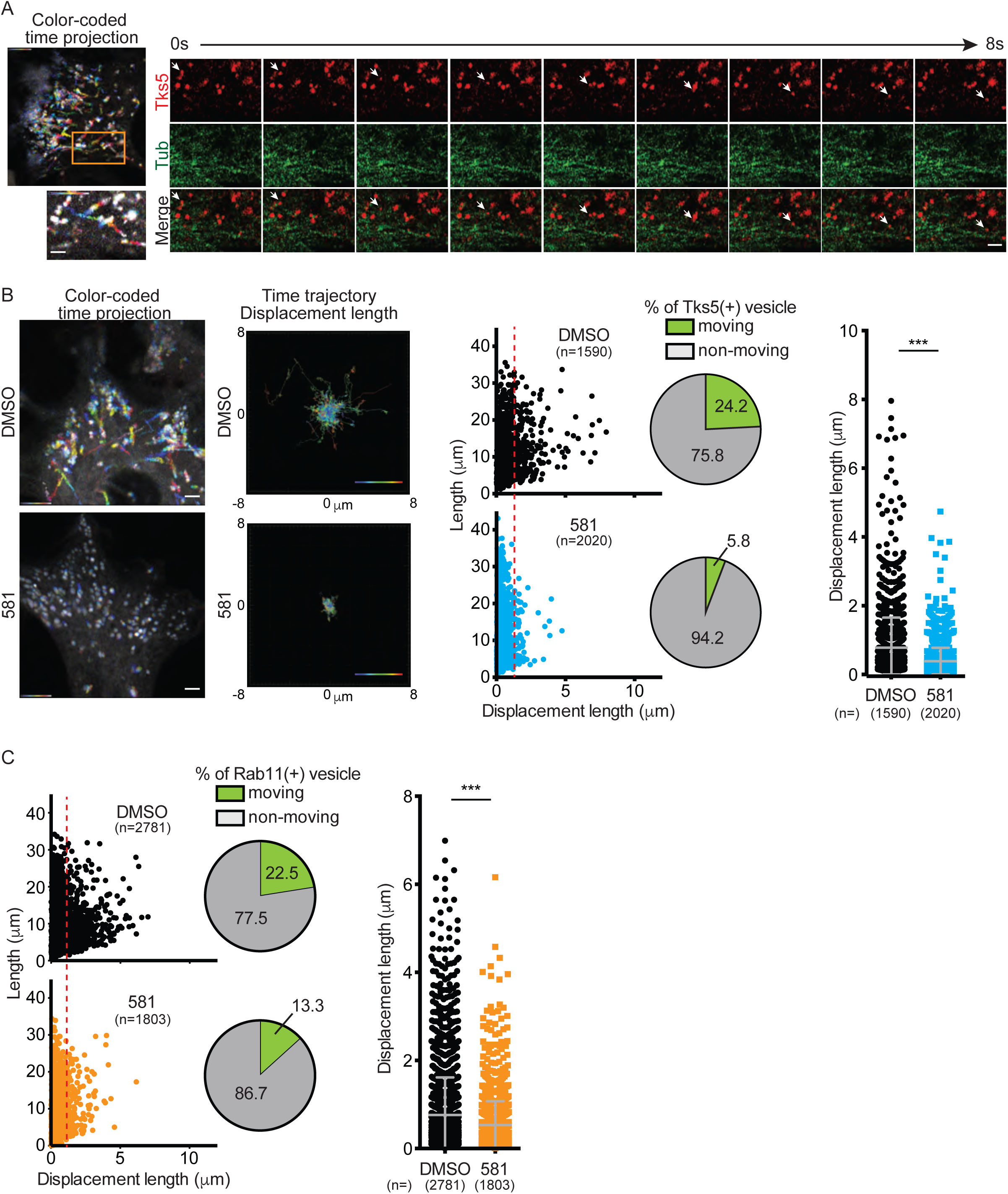
TAO3 regulates trafficking of vesicles containing TKS5. **A** Trafficking of TKS5*α*-positive vesicles was captured by time-lapse imaging (200ms/image for 1 min, total 300 images/film) in Src3T3 cells expressing TKS5*α*-mCherry and YFP-Tubulin. Movie was processed for color-coded time projection (top left) and magnified area (orange square) was shown as color-coded time projection (bottom left) and separated channels in time frame (selected time point during 0-8 sec). **B** SBI-581 inhibits TKS5*α*-positive vesicle trafficking. The movies were taken from cells with treatment of DMSO or SBI-581 (100nM) for 1 min (200ms interval). Trafficking of all TKS5*α*-positive vesicles was shown by color-coded time projection (left), time trajectory displacement length (second left), plotting graph of length/displacement length (middle) with percentage of TKS5*α*+ vesicle moving (pie chart) and displacement length (right). **C** SBI-581 inhibits the trafficking of a fraction of Rab11+ vesicles. The movies were taken from cells with treatment of DMSO or SBI-581 (100nM) for 1 min (200ms interval). Trafficking of all Rab11 positive vesicles was shown by plotting graph of length/displacement length (left) with percentage of Rab11+ vesicle moving (pie chart) and displacement length (right). Scale bars 2 μm (A, B and C). Data shown in panel B and C are mean SEM of biological replicates from 2 or more separated experiments. *P*>0.05 unless other specified; ***, *P*<0.001.

To determine how TAO3 might impact endocytic trafficking of TKS5α, we next initiated a phosphoproteomics screen, using mass spectrometry to identify and compare phosphopeptides in C8161.9 cells in which the endogenous TAO3 was replaced with either wild-type or kinase-inactive TAO3. Table 2 shows the 9 candidates whose phosphorylation was significantly increased in the presence of wild-type TAO3 compared to kinase-dead. Of these, we were particularly interested in cytoplasmic dynein 1 light intermediate chain 2 (LIC2), since dynein complexes act as motors to oppose the actions of kinesin motors to traffic vesicles backwards and forwards along microtubules [43, 44]. In particular, light intermediate chains can recruit the dynein complex to endosomes and lysosomes [45] and also contact adaptors and provide a link to cargo [46]. In keeping with a possible role of LIC2 in TAO3 function, we observed that these two proteins co-localize in the cytoplasm, but not at invadopodia (Figure 6A). We next used knockdown and reconstitution experiments to evaluate the role of LIC2 in invadopodia formation, using both C8161.9 (Figure 6B) and WM793 cells (Supplementary Figure S6A). Knockdown of LIC2 resulted in an increase in invadopodia formation, which was rescued by re-introduction of the wild-type protein. In contrast, introduction of a LIC2 molecule mutated at the identified phosphorylation site (S202A) markedly inhibited invadopodia formation, whereas introduction of a “phosphomimic” form (S202D) promoted invadopodia form to the same extent as LIC2 knockdown. To test whether SBI-581 exerted its effects through preventing TAO3 phosphorylation of LIC2 on S202, we compared the effect of DMSO and SBI-581 on these same cells (Figure 6C and Supplementary Figure S6B). In cells expressing the wild-type protein, SBI-581 inhibited invadopodia formation as expected. However, the inhibitor was without effect in cells lacking LIC2 or expressing the S202A mutant. A small, but significant inhibitory effect was observed in cells expressing S202D LIC2, which may reflect a role for other TAO3 phosphorylation sites on LIC2. Together, these data are consistent with LIC2 acting in an inhibitory fashion in invadopodia formation, and suggest that this inhibition is relieved by phosphorylation of LIC2 on serine 202 by TAO3.

**Figure 6.**
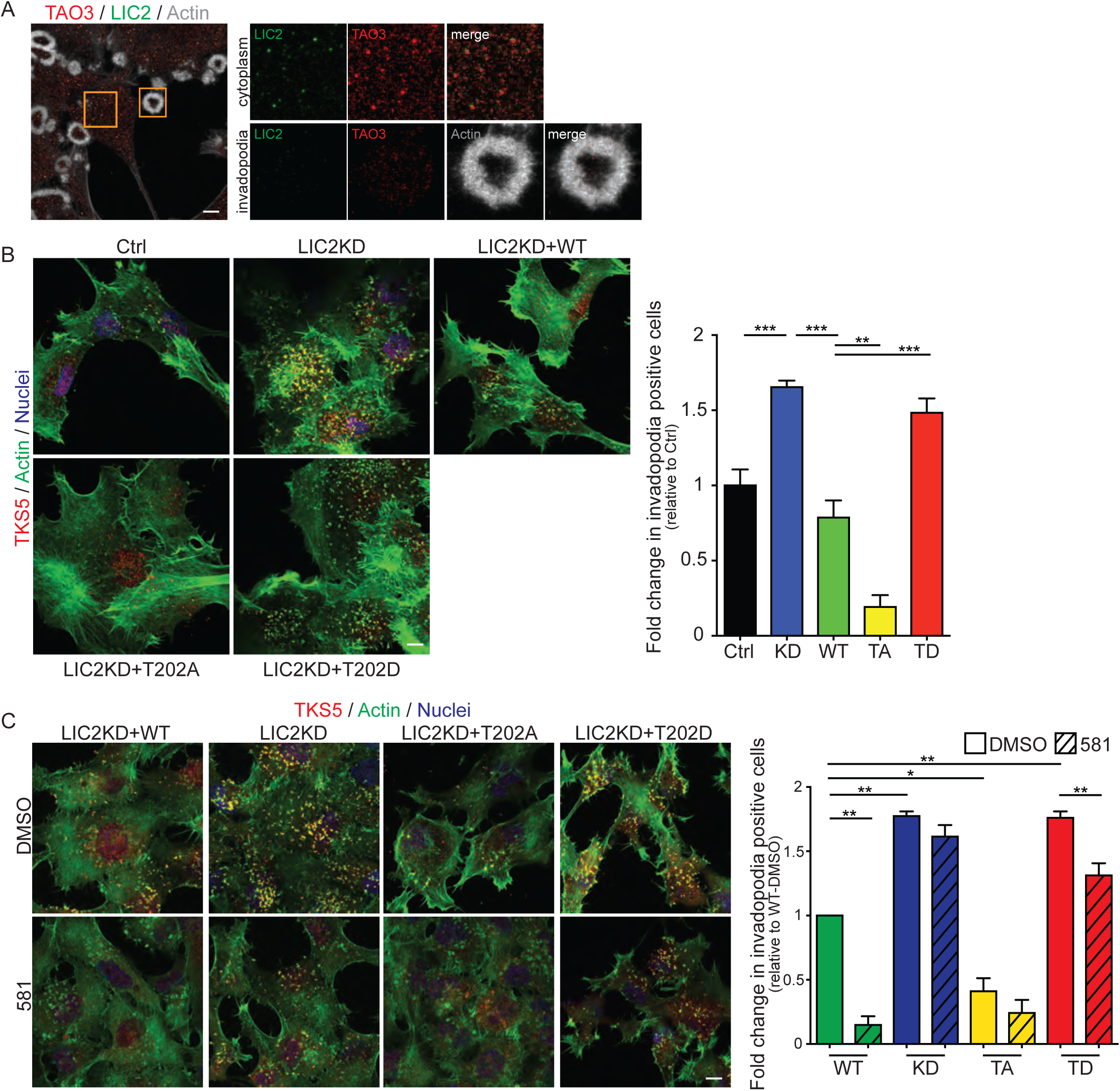
TAO3 phosphorylation of LIC2 promotes invadopodia formation. **A** Colocalization of TAO3 and LIC2. Staining of TAO3 (red), LIC2 (green) and actin (gray) in Src3T3 cells. Images were processed by maximum intensity projection. Magnified area from orange squares (cytoplasm or invadopodia) are shown in right. **B** Invadopodia formation analysis in C8161.9 cells with shRNA-scrambled (Ctrl), shRNA-LIC2 (LIC2KD), shRNA-LIC2+rescued expression of shRNA-resistant LIC2 wild-type (LIC2KD+WT), LIC2-T202A (LIC2KD+T202A) or LIC2-T202D (LIC2KD+T202D). Representative images (left) and fold change in invadopodia positive cells (right). Immunofluorescence staining of invadopodia by actin (phalloidin, green) and TKS5 (red), and Hoechst to denote nuclei of cells. **B** The same Invadopodia formation assay in B with DMSO or SBI-581 (581, 100nM).

**Table 2.** Phosphoproteomics approach to identify TAO3 substrates. *Fold Change: wild-type TAO3 compare to kinase-dead TAO3.

| Protein Name | Peptide Sequence | Phosphorylation site | Fold Change* | P-Value |
| --- | --- | --- | --- | --- |
| Peptidyl-prolyl cis-trans isomerase-like 4 | IPDRS*PEPTR | S178 | 2.77 | 0.00E+00 |
| Utrophin | FEADS*TVIEK | S1025 | 2.41 | 3.58E-09 |
| Zinc finger HIT domain-containing protein 3 | DFLNS*DEEED | S80 | 1.74 | 4.55E-08 |
| ATP-dependent RNA helicase DDX42 | KPVDS*DS*DDDPL | S109: S111 | 1.70 | 1.31E-07 |
| Prostaglandin E synthase 3 | WEDDS*DEDMS | S113 | 1.57 | 3.71E-07 |
| Double-strand break repair protein MRE11 | MANDS*DD SIS | S558 | 1.55 | 1.21E-05 |
| Cytoplasmic dynein 1 light intermediate chain 2 | RGPLT*SGSDE | T202 | 1.54 | 6.77E-06 |
| Sequestosome-1 | EGPSS*LDPSQ | S366 | 1.45 | 1.62E-04 |
| FAS-associated factor 1 | ALRKS*PMMPE | S320 | 1.41 | 4.01E-05 |

## DISCUSSION

Here we used a high-content screening assay to identify kinases required for invadopodia formation, a process that promotes tumor intravasation and extravasation, as well as tumor growth at both primary and metastatic sites. We identified several kinases already implicated in invasive behavior, but we ultimately chose to focus on an “understudied” kinase, TAO3, which we found from public data bases to be robustly expressed in breast cancer, bladder cancer and melanoma, each highly invasive tumors known to elaborate invadopodia. We validated a role for TAO3 in invadopodia formation and function *in vitro*, as well as in extravasation and tumor growth of melanoma *in vivo*. We note that loss of TAO3 expression was not associated with diminished cell viability in our studies, whereas loss of either TAO1 or TAO2 led to rapid cell death. It is important to note that while the N-terminal kinase domains of the three proteins are highly homologous, their C-terminal ∼600-800 amino acid regions are far more distinct from one another. This raises the possibility that the C-terminal regions but not the kinase activity of TAO1 and TAO2 are essential for cell viability. Lastly, in a previous siRNA study using Hela cells, TAOK3 was identified as a survival gene [47]; it is possible that loss of TAO1 or TAO2 (perhaps by RNAi cross-reaction) underlay the observed phenotype in that case. In Drosophila there is a single TAO gene which is involved in both the JNK pathway and actin-microtubule dynamics [48], whereas in more complex organisms, these functions appear to be mediated by different members of the TAO family.

In the course of our experiments, we found that individual knockdown of the 10 kinases from our primary screen showed that they were necessary for invadopodia formation, but 3 of them (AKT3, PKCθ and PYK2) were not required for ECM degradation. To our knowledge, this is the first description of the separation of invasive function from invadopodia formation, and in the future, it will be very interesting to determine the mechanism(s) by which these kinases are involved in invasive behavior.

In order to explore the potential of TAO3 as a drug target, we identified compounds SBI-581 and SBI-029 via high-throughput screening of a kinase inhibitor library and preliminary chemistry. SBI-581 shows high potency against TAO3 (IC_50_=42 nM), moderate selectivity (>5-10x against a broad panel of kinases), and reasonable pharmacokinetics (PK) in mice using IP injection. It therefore represents a good starting point for future optimization. We note, however, that when tested in cells or *in vitro*, SBI-581 inhibited TAO3 function without significant cellular toxicity. This suggests either that some degree of TAO-family specificity has already been achieved with SBI-581 or that the kinase activities of TAO1 and TAO2 are not required for their survival functions. We favor the latter explanation, because we have found that we can rescue the viability of TAO1 knockdown cells by expressing a kinase-dead TAO1 (Supplementary Figure S2B).

We made the unexpected discovery that TAO3 localizes to RAB11+ endosomes situated on microtubules, as well as to invadopodia. Indeed, we also found that RAB11 was required for invadopodia formation, identifying a new link between recycling endocytosis and invasive behavior. Importantly, TAO3 inhibition reduced the movement of recycling endosomes, particularly of those vesicles containing TKS5α, thereby inhibiting invadopodia formation. The regulatory process of endocytosis involves the packaging of selected plasma membrane proteins, and subsequent sorting either for eventual destruction in the lysosome, or recycling back to the membrane [49], with GTPases of the Rab family regulating each discrete step [50]. Receptors, adhesion proteins and extracellular matrix proteins are all targets for endocytosis, and in turn, signaling from these entities can regulate endocytosis. A link between endocytosis and cancer has long been recognized, whereby, for example, oncogenes can promote recycling of key receptors, as well as decrease surface expression of junctional proteins, to promote cell motility [51–53]. Our data expand these properties to include invasive behavior.

We have begun to identify key TAO3 substrates, and validated the dynein protein LIC2 as a target of TAO3 and a negative regulator of invadopodia formation. This suggests a model in which phosphorylation by TAO3 inhibits retrograde vesicular transport, with the net effect of promoting the anterograde transport of TKS5α-containing vesicles to the plasma membrane where invadopodia formation takes place. The mechanism by which TKS5α is loaded onto and off recycling endosomes is obviously of interest, as is the interaction of TAO3 with endosomes. To this latter point, we note that TAO3, kinesin and dynein complexes all contain coiled-coil domains, which are known mediators of both intra- and intermolecular interactions [54]. In a second, unpublished phosphoproteomic analysis, we identified additional TAO3 phosphorylation sites in LIC2, as well as in components of the kinesin complex. In the future, it will be important to define all of the key TAO3 substrates responsible for the phenotype we observe. Nevertheless, the mechanistic underpinnings of TAO3 loss and the potency of the tumor growth and extravasation inhibition we have observed here, coupled with the lack of detrimental effects of TAO3 loss or inhibition in normal cell types, and the high degree of overexpression of TAO3 in several cancer types, supports the preclinical development and eventual clinical testing of selective TAO3 inhibitors.

## ACKNOWLEDGMENTS

We thank Devon Kelly for assistance in identification and acquisition of human tumor samples from the OHSU Biolibrary, the OHSU Histopathology and Proteomics Shared Resources for their support, Susanne Heynen-Genel (SBP) for her support in the establishment of the high content screening assay, and Pam Carroll for spearheading the generation of the mouse kinome siRNA library. We acknowledge the expert technical assistance by Stefanie Kaech Petrie and Crystal Chaw in the Advanced Multiscale Microscopy Shared Resource (NIH P30 CA069533), and Larry David, Jennifer Cunliffe, John Klimek, and Phillip Wilmarth in the Proteomics Shared Resource at the OHSU Knight Cancer Institute. We thank the personnel and support from the Cancer Center Structural Biology Core at the Sanford Burnham Prebys Medical Discovery Institute (SBP), and at the Lentivirus Production Cores at SBP and OHSU.

## METHODS

### Cell lines

The C8161.9 human melanoma cell line was purchased from ATCC. The WM793 human melanoma cell line was a gift from Dr. Gary G. Chiang (Abbvie, Chicago, IL) and authenticated by OHSU DNA core facility using short tandem repeat analysis. Bladder cancer cell lines, T24 and UM-UC-3 and breast cancer cell line, Hs578t were purchased from ATCC. Luciferase-expressing human breast cancer cell line MDA-MB-231-Luc was obtained from Xenogen. The Src-transformed NIH-3T3 (Src3T3) cells have been described before in [55]. Melanoma, breast cancer cell lines and Src3T3 cells were grown in Dulbecco’s Modified Eagle’s Medium (10-013-CV, Corning) containing 10% fetal bovine serum (FBS, HyClone and Gibco). T24 was cultured by McCoy’s 5a Medium Modified (ATCC) with 10% FBS. UM-UC-2 was grown in Eagle’s Minimum Essential Medium (ATCC) with 10% FBS. All cell lines were routinely tested for mycoplasma contamination and confirmed negative for mycoplasma species.

### Antibodies and reagents for staining

Antibodies used for immunoblotting were: TAO3 (ab150388, 1:1000, Abcam), CAMK1 (ab68234, 1:1000, Abcam), RAB11 (610657, 1:1000, BD Biosciences) and tubulin (T6557, 1:3000, Sigma). Antibodies used for immunofluorescence staining were: TAO3 (LS-B11301, 1:250, LSBio), tubulin (6074, 1:250, Sigma) and RAB11 (71-5300, 1:250, Thermo Fisher Scientific). The antibodies for TKS5 (Rabbit monoclonal, F4 and mouse monoclonal, G6) were generated and validated by the Courtneidge laboratory, and will be fully described in a separate publication (in submission). Phalloidin (Alexa fluor-350, −488, −568 or −647, Thermo Fisher Scientific) was used for actin staining. For the gelatin degradation assay, Oregon green gelatin (0.2 mg/ml in PBS containing 2% sucrose; Invitrogen) was used. Hoechst (1:4000, Thermo Fisher Scientific) was used for nuclear staining.

### Screening and validation

The high content screening assay for invadopodia formation has been previously described [24]. Briefly, we ran the screen twice, each time in quadruplicate, with a negative control (non-target scrambled siRNA). We used pools of three different siRNAs for each gene. The results from the two runs had a low coefficient of variability, therefore we did not repeat the assay a third time. The effects of siRNA knockdown on invadopodia number were determined by eye, counting at least 100 cells for each field. We considered cells as positive if they had more than two invadopodia. siRNAs that triggered invadopodia inhibition < 50% (compared to control) in cells that retained viability and otherwise contained a normal actin cytoskeleton were considered hits. In order to validate the screening results, we used the three individual siRNAs (Mission siRNA, Sigma). The validation step was repeated three times and we scored for phenotypic concordance. For each hit, we tested the effect of kinase knockdown on gelatin degradation (using the pool siRNA only). The siRNA oligo IDs are listed in Supplementary Table S5.

### Expression analysis

Cancer Outlier Profile Analysis (COPA) was performed on public gene expression data sets curated by Oncomine [32]. Gene expression values were median-centered, setting each gene’s median expression value to zero. The median absolute deviation (MAD) was calculated and scaled to 1 by dividing each gene expression value by its MAD. The 75th, 90th, and 95th percentiles of the transformed expression values were calculated for each gene, and then genes rank-ordered by their percentile scores, providing a prioritized list of outlier profiles. A gene rank over-expression threshold of 10% was set for this analysis For each gene identified as an outlier in at least one dataset, we tabulated all data sets with outlier calls and the total number of datasets for that cancer type with expression of that gene.

### Immunoblotting

Cell lysates were prepared by washing cells twice with cold Tris-buffered saline (TBS) containing 100μM Na3VO4 and then lysing in 50 mM Tris-HCl (pH 7.5), 250mM NaCl, 1% Triton X-100, 50 mM NaF, 100μM Na3VO4 and 1mM EDTA lysis buffer containing a dissolved complete mini protease inhibitor tab (Roche Diagnostics, Germany). Supernatant of cell lysates was assayed for total protein content using the BCA protein assay (Thermo Fisher Scientific, Rockford, IL), and 70 μg of total protein per sample was separated in a 7.5% polyacrylamide gel (Invitrogen). Secondary antibodies were conjugated to Alexa Fluor 680 or IR800, and membranes were scanned using an infrared imaging system (Odyssey; LI-COR Biosciences, Lincoln, NE).

### RT-qPCR

RNA was isolated from cell lines using the RNeasy RNA isolation kit (Qiagen). cDNA was generated using the SuperScript III First-Strand Synthesis kit (Invitrogen). Absolute expression analysis was done by using standard curve for each primer sets that made by DNA plasmids containing each isoforms as template, and the gene copy number was measured as copies per 1μg RNA input. The following primer sets were used:

hCamk1-F: TACAGCAAGGCTGTGGATTG

hCamk1-R: AGTGCCGGATGAAATCTTTG

hPak6-F: AATGGCAGAACATCCTGGAC

hPak6-R: TTCTGGATGTCGTTGAGCAG

hTao3-F: AGGACCATAGCACACCCAAG

hTao3-R: AATGTTCTGCTGCTCCACCT

### DNA constructs and siRNAs

siRNA oligos targeting human TAO1 (pool; M004846-08), TAO2 (pool; M004171-03), TAO3 (pool; M004844-02), PAK6 (pool; M004338-02), CAMK1 (pool; M004940-00) and RAB11 (pool; set of 4, LQ-004726-00, two individual; J-004726-07 and J-004726-09) as well as non-targeting controls were obtained from Thermo Fisher Scientific.

pGIPz lentiviral shRNAs used for non-silencing control, human TAO3 or LIC2 knockdown were purchased from Dharmacon or Millipore Sigma. The clone used were V3LMM_418096 (TAO3) and TRCN0000116993 (LIC2). Lentiviral plasmid pCDH-CMV-MCS-EF1-Puro (Addgene) was used for overexpression of human TAO3-SR (shRNA resistant), human TAO3-KR (shRNA resistant and kinase dead), human TKS5α-mCherry, YFP-Tubulin, Lifeact-mCherry, human LIC2-WT (shRNA resistant), human LIC2-T202A and human LIC2-T202D. Plasmids encoding wild-type human RAB11 (#12679) was obtained from Addgene. The following primer sets were used for site directed mutagenesis:

TAO3-SR-F: 5’-caggacacttgcaaagtgcaaacgaaacagtataaagcac-3’,

TAO3-SR-R: 5’-gtgctttatactgtttcgtttgcactttgcaagtgtcctg-3’.

TAO3-KR-F: 5’-gaggtggtggcaattaggaagatgtcctatagtggg-3’,

TAO3-KR-R: 5’-cccactataggacatcttcctaattgccaccacctc-3’,

LIC2-shRNA resistant-F: 5’-gaaaaacctcgacttgttatacaaatatattgttcataaaac-3’,

LIC2-shRNA resistant-R: 5’-gttttatgaacaatatatttgtataacaagtcgaggtttttc-3’,

LIC2-T202A-F: 5’-cggagcctgaggccagagggcctct

LIC2-T202A-R: 5’-agaggccctctggcctcaggctccg-3’,

LIC2-T202D-F: 5’-ttcatcggagcctgagtccagagggcctcttctc-3’,

LIC2-T202D-R: 5’-gagaagaggccctctggactcaggctccgatgaa-3’.

Lentiviral preparations were made by the viral core facilities at the Sanford|Burnham|Prebys Medical Discovery Institute (La Jolla, CA) and the Oregon National Primate Research center (Molecular & Cell Biology, Lentivirus Service) at the Oregon Health and Science University (Portland, OR).

### Invadopodia assay and degradation assay

Invadopodia staining and gelatin degradation assays were performed as previously described [7]. Briefly, cells were grown on glass coverslips with or without collagen (invadopodia formation assay) or gelatin-coated coverslips (degradation assay) and fixed with 4% paraformaldehyde/PBS (Electron Microscopy Sciences). For the invadopodia assay on collagen-coated coverslips, high-dense fibrillar type I collagen (HDFC) was prepared according to original protocol reported [56, 57]. Briefly, 18mm coverslips were pre-chilled on ice and coated with 10μl ice-cold neutralized collagen (#35429, Corning). The pipette tip was used to spread the collagen evenly on the glass surface and the coated coverslips were left on ice for 10 min to facilitate flattening of the collagen. The layer of collagen was polymerized into a fibrillar meshwork at 37°C for 30 min, followed by centrifuged at 3,500 g for 20 min. After fixation and permeabilization with 0.1% Triton X-100/PBS for 15 min, the cells were blocked by 5% BSA in PBS with 5% goat-serum for 1 hour at room temperature (RT) and incubated with primary antibodies for 90 min at RT (or overnight at 4°C). The cells were washed and incubated with Alexa Fluor-conjugated secondary antibodies and phalloidin. Fluorescence microscopy images were obtained with a fluorescent microscope (Axioplan2; Carl Zeiss) equipped with a charge-coupled device camera (AxioCam HRm; Carl Zeiss). For each invadopodia experiment, the number of cells forming invadopodia was quantified in ≧5 microscope fields (63x) imaged randomly and percentage of invadopodia forming cells was assessed. For each ECM degradation experiment, ≧6 microscope fields (40x) were imaged randomly. The percentage of degraded area was quantified with ImageJ software (National Institutes of Health) and normalized to the number of nuclei in that area was represented as “% degradation per cell”.

### High-resolution invadopodia and time-lapse imaging

Src3T3 cells were cultured on glass coverslips and prepared for imaging using the same methodology as for the invadopodia assay. Confocal images were collected using a laser-scanning confocal microscope LSM880 equipped with AiryScan (Carl Zeiss). Time-lapse imaging was performed by 200ms per frame for 1 min, total 300 frames. Images were transferred to Imaris™ (Bitplane) which is a multidimensional analysis program based on the fluorescence intensity data. For colocalization analysis, surfaces (spots) were created on the TKS5, TAO3 and RAB11 separately and colocalized signals were assessed by “ImarisColoc” tool. For vesicle trafficking, surfaces (spots) were created on the TKS5α-mCherry or RAB11-DsRed to track the vesicles and extract the “displacement length (μm)” and “Length (μm)” as statistics Excel files. For making movies, the time-lapse films were edited by Apple’s iMovie software.

### IHC and scoring

Primary melanomas were stained with TAO3 (ab150388, 1:300, Abcam) on a Ventana Discovery autostainer. Negative control slides were stained with Rabbit IgG. Subsequently, TAO3 IHC was scored by a board-certified pathologist (G.V.T). Scores were either 0 for no staining in the tumor, 1+ for weak staining, 2+ for moderate to strong staining. Next, an immunoscore was calculated from the formula: (0 x % cells staining negative) + (1 x % cells staining weakly positive) + (2 x % cells staining moderately-strongly positive), giving a range of 0-200.

### Invasion assays

Boyden chamber invasion assays were performed as previously described [55]. Briefly, Matrigel-containing transwell chambers (#08-774-122, Corning) were pre-hydrated overnight. The cells with medium containing 0.1% FBS were placed in the insert chambers and media containing 10% FBS was placed in the lower chambers to facilitate chemotaxis. Invasion assays were run for 16 hr. Noninvading cells were removed from top side of the insert chamber, then the images of GFP+ cells which passed through the matrigel membrane were taken by fluorescence stereo zoom microscope (Zeiss Axio Zoom). Invaded cells were assessed by GFP intensity using ImageJ software.

### 2D/3D proliferation assay

Type I collagen 3D cultures were performed as described previously [7]. Briefly, rat tail type I collagen (#354263, Corning) was prepared to a final concentration of 2.1 mg/ml, and 5,000 to 25,000 cells were added to the collagen mix before gelling. Spread cells were grown for 12 days in DMEM containing 10% FBS. The matrix was dissolved with 2 mg/ml collagenase type2 (#LS004176, Worthington Biochemical corporation) and cell numbers were determined by hemocytometry. Type I collagen 2D cultures were performed according to the manufacturer’s instructions. Briefly, 50 μg/ml rat type I collagen in 0.02M acetic acid was incubated with coverslips at RT for 1hr. After washing, cells were added and cultured for 8 days (C8161.9) or 9 days (WM793) in DMEM containing 10% FBS, and cells were counted. The proliferation assay on plastic plates was performed by the same method described previously without type I collagen coating.

### Spheroid invasion and growth assay

3D spheroid cultures were performed as described previously [58]. Briefly, spheroids of C8161.9 cells were prepared in hanging droplets with 5,000 cells in 10μl of 20% FBS containing DMEM for 5 days. The spheroids were embedded in 2.1 mg/ml of type I collagen and incubated with DMEM containing 10% FBS for 2 days. Spheroids were fixed by 4% PFA and washed by PBS for three times, then stained by Alexa568-phalloidin to visualized overall of spheroid in 3D collagen. Imaging was performed on Zeiss Axio Imager 2 equipped with ApoTome.2 (Zeiss) and the z-section with maximum size of actin stained spheroids were taken and measured the “spheroid size (actin intensity)”.

### Tumor growth in mice

All animal experiments were conducted in accordance with the NIH Guide for the Care and Use of Laboratory Animals. Subcutaneous implantation was performed with minor modifications as described previously [7]. Briefly, cells were resuspended in PBS/Matrigel mixture (1:1 ratio, BD Biosciences) to a final concentration of 4x 10^6^ cells/ml. Athymic nude mice (nude Nu/J, The Jackson Laboratory) were injected in the flank with 100 μl, and tumors were allowed to form for 22 days. For the tumor growth experiment with inhibitor, SBI-581, tumors were allowed to form for 10 days and SBI-581 (final concentration 10 mg/kg in DMSO with 1:5 dilution of PBS) were injected every day for 10 days. Tumor growth was measured every 2-3 days using calipers. The longest (L) and shortest (S) measurements were recorded and tumor volumes were calculated as Volume = 0.5*(L x S2) and expressed as mean volume (mm2).

### Extravasation efficiency assay in mice

Extravasation efficiency assay was performed as described previously [17]. GFP expressing C8161.9 cells with pLKO.1-scrambled, TAO3 knockdown (KD) or rescued (SR) were injected intravenously in tail veins of 8 week old SCID/Beige female mice (C.B-17/IcrHsd-Prkdc scid Lyst bg-J, ENVIGO). 24 hours after injection of 2.5 x 105 of cells, mice were euthanized and tumor cells in lung blood vessels were removed by perfusion, then lungs were harvested. Frozen serial sections of all lungs (50 μm thickness) were taken every five sections. For the efficiency analysis, slides were counter stained with Hoechst for 15 min before mounting. All sections were scanned by Zeiss AxiaScan system with ZEN Blue software. GFP positive cells were counted from all scanned lung sections then calculated as “Number of GFP+ cancer cells in lung”. For inhibitor experiment, DMSO control or SBI-581 (30mg/kg) was injected intraperitoneally, then 30 minutes later GFP-labeled C8161.9 cells were injected through tail vein.

### Phosphoproteomic analysis

Grow C8161.9 cells with endogenous TAO3 replacement by shRNA-TAO3 and wild-type TAO3 (shRNA-resistant, SR) or kinase-dead TAO3 (shRNA-resistant, KR) on five 150 mm culture dishes grown to between 70-80% confluence. Cells were washed with 1X cold-PBS before lysis to remove any media containing protein contaminants. Lyse the cells by 10 ml of Urea Lysis Buffer (at room temperature, 50 mM HEPES pH 8.5, 8 M urea, 1 mM NaF, 1 mM sodium orthovanadate, 10 mM sodium β-pyrophosphate, 1 mM β-glycerophosphate buffer) and disrupted by sonication on ice. The cell lysates were stored at −80°C prior to digestion and enrichment. Protein concentrations were then determined using the Pierce™ BCA Protein Assay Kit (Thermo Scientific). Approximately 12 mg of protein from each lysate was used. Protein was reduced with dithiothreitol, alkylated with iodoacetamide, urea diluted to 2 M, then trypsin (TPCK treated, Worthington) was added at a 25:1 protein:trypsin ratio. The samples were incubated overnight at 37°C before being quenched with TFA at a final concentration of 1%. The peptides were then solid phase extracted using Waters Sep tC18 cartridges (Waters Corporation). Peptide concentrations were then determined using the Pierce™ Quantitative Colorimetric Peptide Assay (Thermo Scientific) and dried by vacuum concentration. The phosphopeptides were enriched using previously published methods [59, 60] using Titanosphere TiO_2_ 5 μm particles (GL Biosciences). The enriched phosphopeptides were purified by solid phase extraction using UltraMicroSpin columns (The Nest Group, Inc.) and dried down in preparation for TMT labeling. The enriched phosphopeptides from the SR and KR cells were then labeled with TMT reagents as recommended by the manufacturer (Thermo Scientific), mixed, and analyzed by two-dimensional liquid chromatography/mass spectrometry using an Orbitrap Fusion Tribrid mass spectrometer (Thermo Scientific) as previously described [61]. The full proteomic dataset can be found in the PRIDE database with the following accession number (provided upon publication).

### Protein production

The protein (TAO3 kinase domain (1-319) fused with GST) was expressed in the baculovirus intracellular expression system. Linear DNA was used for the transfection to make virus. Protein was expressed using 8 liters of SF9 cells. After 72 hours of infection, cells were collected by centrifugation, re-dissolved in a buffer containing 50 mM NaCl, 20 mM TRIS pH 8, 1 mM EDTA, 2 mM of β-mercaptoethanol and including a standard protease inhibitor cocktail. Cells were lysed in this buffer with a glass dounce homogenizer on ice. The supernatant was separated from pellet by centrifugation for 1 h at 20K rpm (SS34 rotor) and then incubated with glutathione–agarose bead for 2 HRS at 4°C. Beads were washed with 50 mM NaCl, 20 mM TRIS pH 8, 1 mM EDTA, 5 mM β-ME and then the TAO3 kinase domain was cleaved off the beads by incubation with 3C protease. Cleaved released kinase domain was separated from contaminating GST by immediately running the sample in this buffer onto a mono-Q on the AKTA FPLC, eluting protein with a gradient of the same buffer containing 50 to 1 M NaCl. The peak containing the TAO3 kinase domain was collected and then run on a Superdex S200 gel filtration column prequilibrated with the final buffer 100 mM NaCl, 25 mM Tris-Cl pH 8.0, 2 mM β-ME, 0.3 mM EDTA.

### High throughput screen for TAO3 Inhibitors

A time-resolved fluorescence resonance energy transfer (TR-FRET) assay was developed to monitor the inhibition of TAO3 kinase activity. We used the kinase domain of TAO3, and an HTRF-KinEASE assay along with the S3 substrate and a Eu^3+^ Cryptate-conjugated anti-pSer/Thr antibody. This assay was miniaturized to 1536 well format with a final volume of 2µL + 2µL detection reagent. Compounds (including a known kinase inhibitor collection (800 compounds) and an SBP Institute-selected kinase inhibitor scaffold library (4,800 compounds)) were tested at 10µM. Assay performance was very robust with an average Z’ of 0.89, and with no individual plate with a Z’ of less than 0.70. Hit threshold at 30% inhibition, and since the hit rate was low (not unexpected based on the lack of reported TAO3 inhibitors in the scientific and patent literature) this resulted in 82 hits (0.12%). To further prioritize confirmed hits, apparent IC_50_ values were determined. Twelve selected compounds with potencies <2µM were validated using Protein Thermal Shift (PTS) technology, which demonstrated direct binding of the compounds to TAO3. These compounds were also tested in dose response against an unrelated kinase (MEKK3) using the same assay format, with a minimum selectivity threshold of 5x. From these efforts a series of oxindoles were identified as the most promising hits. Preliminary chemistry led to SBI-581 (IC_50_= 42 nM against TAO3, IC_50_=237 nM against MEKK3). Profiling against the Reaction Biology Corporation KinaseScan panel of 363 kinases and displayed moderate promiscuity with sub-μM activity against approximately 40 kinases.

### Synthesis of SBI-581 and SBI-029

All reactions involving air and moisture-sensitive reagents and solvents were performed under a nitrogen atmosphere using standard chemical techniques. Anhydrous solvents were purchased and freshly used from Sigma-Aldrich or EMD Biosciences. All organic reagents were used as purchased. Analytical thin-layer chromatography was performed on Partisil K6F silica gel 60 Å, 250 μm. ^1^H and ^13^C chemical shifts are reported in δ values in ppm in the corresponding solvent. All solvents used for chromatography on the synthetic materials were Fisher Scientific HPLC grade, and the water was Millipore Milli-Q PP filtered. LCMS analysis of synthetic materials was completed on a Waters Autopurification system, which consists of a 2767 sample manager, a 2545 binary gradient module, a system fluidics organizer, a 2489 UV/vis detector, and a 3100 mass detector, all controlled with MassLynx software. A Sunfire Analytical C18 5 μm column (4.6 × 50 mm) and stepwise gradient [10% (MeCN + 0.1% TFA) in (water + 0.1% TFA) to 98% (MeCN + 0.1% TFA) in (water + 0.1% TFA) for 9 min.] was used for analytical LCMS of final compounds. The final compounds were purified as specified. All NMR spectra for the synthetic materials were recorded on a Bruker Avance II 400 MHz instrument. The MestReNova 7 program was used to process and interpret NMR spectra.

#### Synthesis of SBI-581

Step 1: Preparation of (Z)-3-((1H-indol-2-yl)methylene)-4-iodoindolin-2-one: To a solution of 4-iodoindolin-2-one (1.5 g, 5.8 mmol) in ethanol (30 mL) was added piperidine (49.2 mg, 0.58 mmol) and 1H-indole-2-carbaldehyde (1.45 g, 11.6 mmol). The mixture was refluxed overnight. The reaction was cooled to room temperature and the resulting orange solid was filtered and washed with ethanol and dried. The orange solid (1.15 g, Yield 93 %, 95 % pure by HPLC) (Z)-3-((1H-indol-2-yl)methylene)-4-iodoindolin-2-one was used as in the next reaction. MS: m/z 387.1.

Step 2: Preparation of tert-butyl (Z)-4-((3-((1H-indol-2-yl)methylene)-2-oxoindolin-4-yl)ethynyl)-4-hydroxypiperidine-1-carboxylate: To a solution of (Z)-3-((1H-indol-2-yl)methylene)-4-iodoindolin-2-one (1.15 g, 2.97 mmol) in DMF (20 mL) was added tert-butyl 4-ethynyl-4-hydroxypiperidine-1-carboxylate (662 mg, 2.97 mmol), Pd(PPh_3_)_4_ (200 mg, 0.17 mmol), CuI (200 mg, 1.05 mmol) and triethylamine (624 mg, 6.12 mmol). The mixture was carefully degassed with N_2_. The mixture was then refluxed overnight under N_2_. The reaction was then evaporated to remove the DMF. The remaining mixture was diluted with water (30 mL) and extracted with ethyl acetate (50 mL). The organic layer was concentrated *in vacuum* and the residue was purified by flash column chromatography over silica gel (ethyl acetate/hexane gradient: 0 to 80%) to afford tert-butyl (Z)-4-((3-((1H-indol-2-yl)methylene)-2-oxoindolin-4-yl)ethynyl)-4-hydroxypiperidine-1-carboxylate (620 mg, yield: 54%) as a red solid. MS: m/z 484.6.

Step 3: Preparation of (Z)-3-((1H-indol-2-yl)methylene)-4-((4-hydroxypiperidin-4-yl)ethynyl) indolin-2-one (SBI-581): To a solution of tert-butyl (Z)-4-((3-((1H-indol-2-yl)methylene)-2-oxoindolin-4-yl)ethynyl)-4-hydroxypiperidine-1-carboxylate (620 mg) in 30 ml of dry methylene chloride at 0°C was added 5 ml of TFA. The reaction was stirred and slowly warmed to RT. The progress of the reaction was monitored by HPLC. When the reaction was complete (∼2 hr), the solvent was removed under vacuum and the residue was directly purified by preparative HPLC (C_18_-silica, MeCN in water: 5% to 95% gradient) to afford (Z)-3-((1H-indol-2-yl)methylene)-4-((4-hydroxypiperidin-4-yl)ethynyl)indolin-2-one (90 mg, yield: 19%) as a red solid (>95% purity). ^1^H NMR (400 MHz, DMSO-*d*6): δ 8.76(1H, s), 7.65 (1H, 2H, J = 3.2 Hz), 7.61 (1H, d, J = 3.2 Hz), 7.61 (1H, d, J = 3.0 Hz), 7.28 (1H, s), 7.24 (2H, 2H), 7.12 (3H, s. m), 6.97 (1H, d, J = 3 Hz), 3.2-2.9 (4H, bm), 2.02 – 2.18 (3H, bm), 1.82 – 1.95 (2H, bm); MS: m/z 384.5

#### Synthesis of SBI-029

SBI-029 was synthesized in a similar fashion to SBI-581 using 5-fluoro-4-iodoindolin-2-one and 4-methoxy-1*H*-pyrrole-2-carbaldehyde as starting materials.

### Rodent pharmacokinetics of SBI-581

SBI-581 was dosed at 1 mg/kg iv (as a 1 mg/mL solution in 75%PEG300/25% water), 10 mg/kg po (as a 3 mg/mL solution in Pharmatek #6) and 10 mg/kg ip (as a 3 mg/mL solution in 5% DMSO/5% Tween 80/90% water) to fasted male C57BL/6 mice (3 mice per cohort). Plasma samples were taken at 0.083, 0.25, 0.5, 1, 2, 4, 6 and 24 hr post dose (iv) and 0.25, 0.5, 1, 2, 4, 6 and 24 hr post dose (po and ip) and measured for the level of SBI-581 via LC/MS/MS analysis.

### Quantification and statistical analysis

The numbers of samples (technical replicate) and times of experiments (biological replicate) are indicated in each figure or figure legend. All data points and error bars represent the means and SEMs. GraphPad Prism software was used to make graph and calculate significance by Student’s *t* test. *P* value of <0.05 was considered to be statistically significant and indicated in figure legends.

---

**Suuplementary Figure S1.**
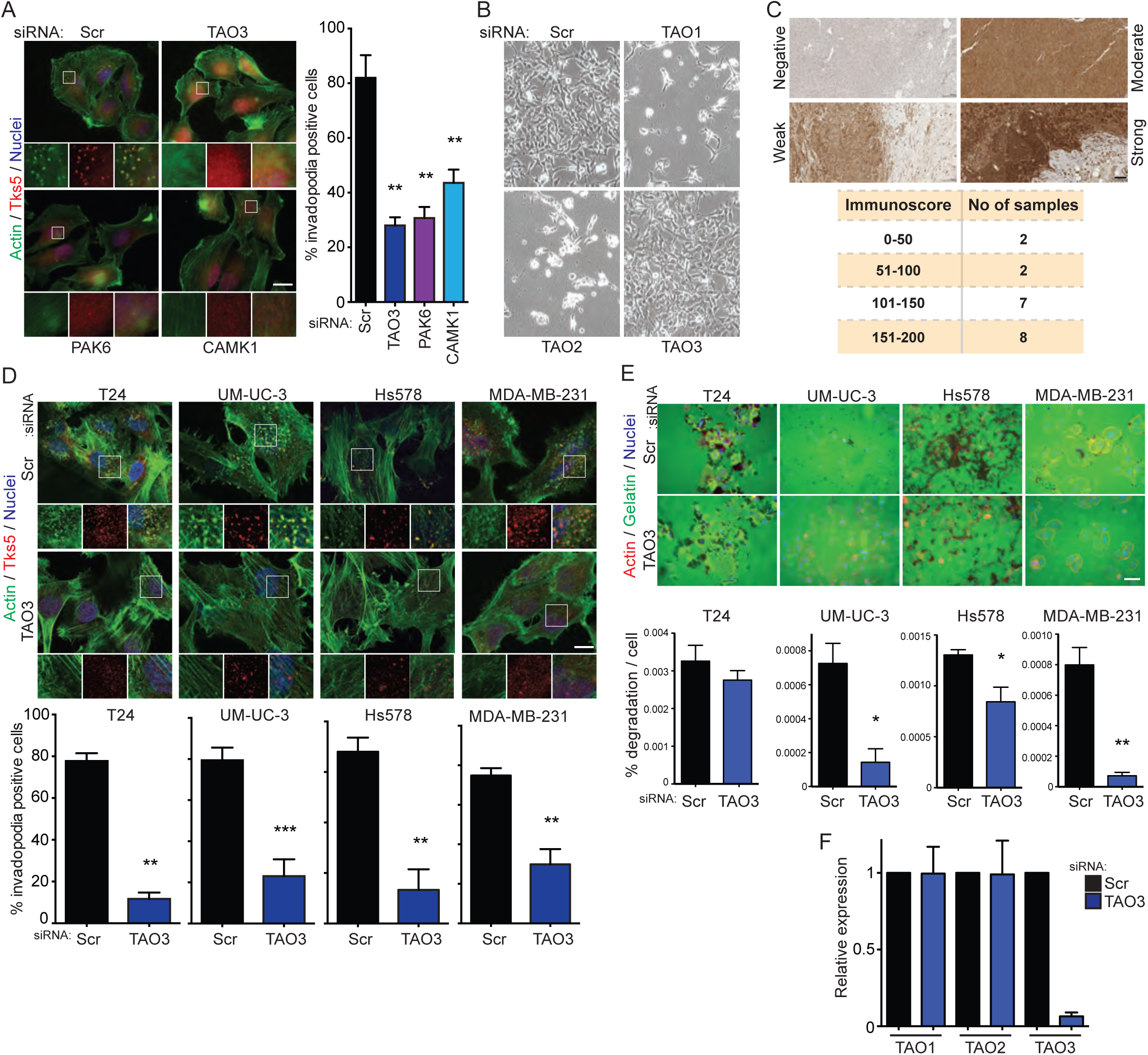
A, Invadopodia formation analysis in WM793 cells with siRNA-scrambled, -TAO3, -PAK6 or –CAMK1. Representative images (left) and percentage of invadopodia positive cells (right). Immunofluorescence staining of invadopodia by actin (phalloidin, green) and Tks5 (red), and Hoechst to denote nuclei of cells. B, Representative bright-field images of C8161.9 for 3 days after transfection with siRNA-scrambled, -TAO1, -TAO2 or -TAO3 in C8161.9. C, Representative IHC staining of TAO3 in melanoma cases (left) and immunoscore (right). Invadopodia formation (D) and gelatin degradation (E) analysis in bladder cancer (T24 and UM-UC-3) and breast cancer (Hs578t and MDA-MB-231) cell lines with siRNA-scrambled or -TAO3. Representative images (top) and percentage of invadopodia positive cells or percentage of degradation per cells (bottom). D, Immunofluorescence staining of invadopodia by actin (phalloidin, green) and Tks5 (red). E, Immunofluorescence staining of actin (phalloidin, red) and gelatin (green). Hoechst to denote nuclei of cells. F, Relative expression of TAO1, TAO2 and TAO3 in C8161.9 cells with siRNA-scrambled and -TAO3. Scale bars, 20 μm (A and D) 50 μm (E) and 100 μm (C). Data shown are n=3 to 10 in each experimental group and were validated in 2 separated experiments. *P*>0.05 unless other specified; *, *P*<0.05; **, *P*<0.01; ***, *P*<0.001.

**Supplementary Figure S2.**
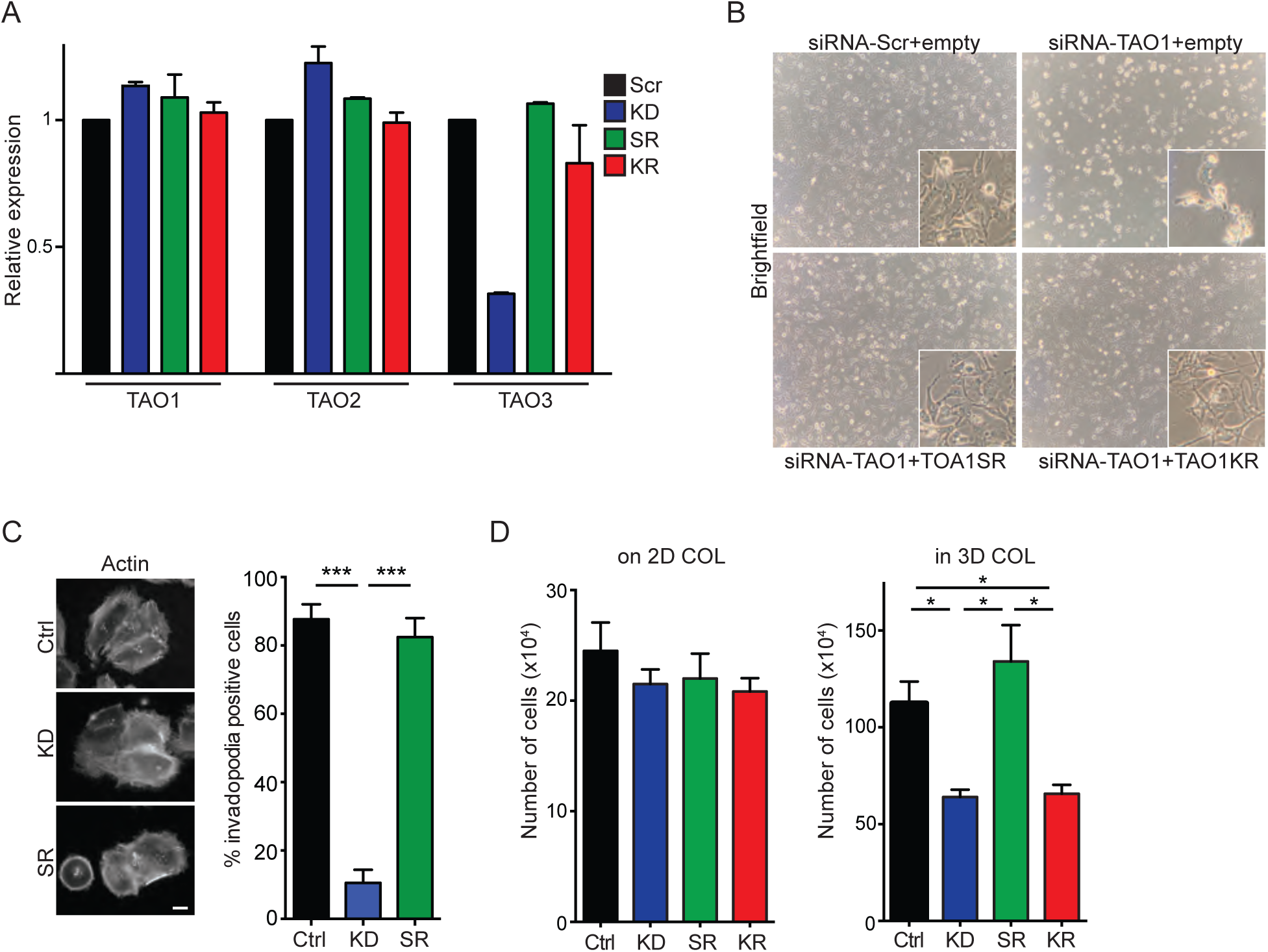
A, Relative expression of TAO1, TAO2 and TAO3 in C8161.9 cells with shR-NA-scrambled (Scr), shRNA-TAO3 (KD), shRNA-TAO3+rescued expression of shRNA-resistant TAO3 (SR) or shRNA-TAO3+rescued expression of shRNA-resistant kinase-dead TAO3 (KR).B, Representatie brightfield image of C8161.9 cells with siRNA-scrambled or siRNA-TAO1, and rescued expression of shRNA-resistant-TAO1 (SR) or shRNA-resistant kinase-dead TAO1 (KR). C, D, TAO3 regulates invadopodia formation and function in melanoma cell line, WM793. Invadopodia formation (C) and 3D growth analysis (D) in WM793 cells with Scr, KD, SR or KR. C, Representative images (left) and percentage of invadopodia positive cells (right). Immunofluorescence staining of invadopodia by actin (phalloidin, green) and Hoechst to denote nuclei of cells. D, Growth of cells are indicated in the figure on 2D type I collagen (on 2D COL, day 9) and in 3D type I collagen (in 3D COL, day 12). Data shown are n=3 and were validated in 2 separated experiments. *P*>0.05 unless other specified; *, *P*<0.05; ***, *P*<0.001.

**Supplementary Figure S3.**
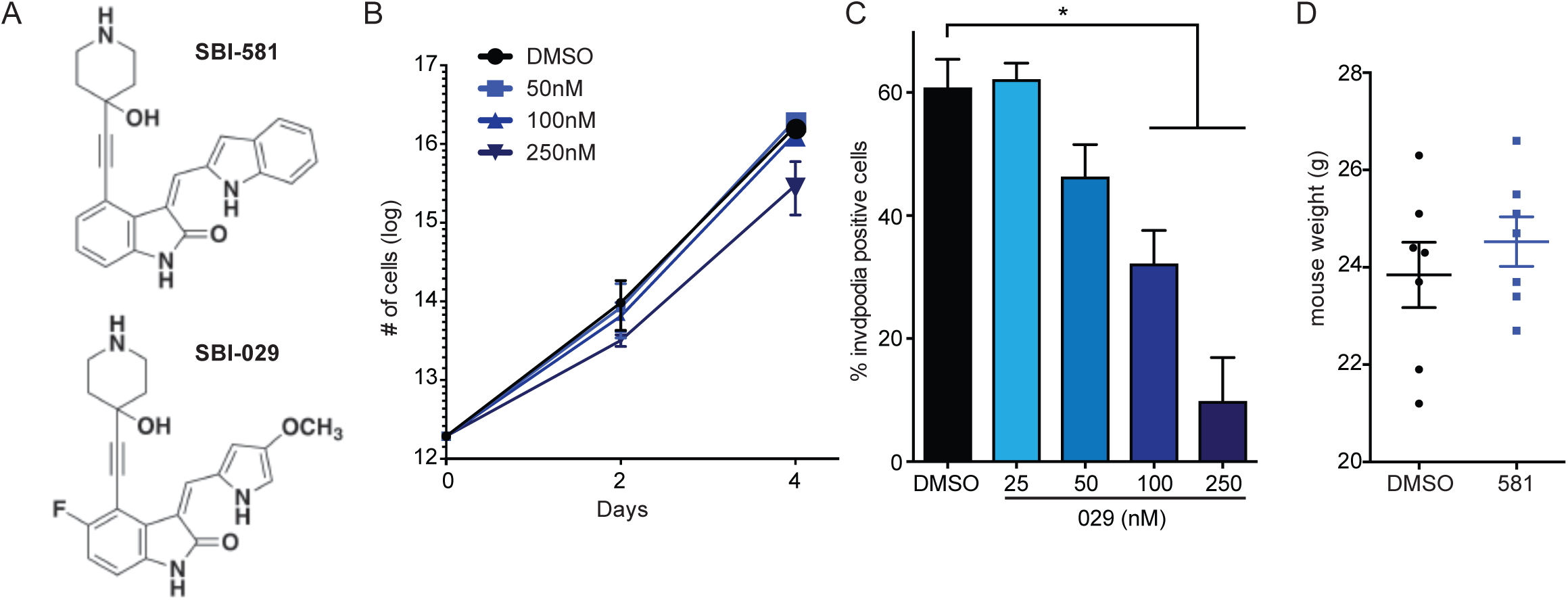
A, Chemical (2D) structure of the TAO3 inhibotors, SBI-581 and SBI-029. B, Cell proliferation of C8161.9 cells on 2D plastic with control DMSO and TAO3 inhibitor, SBI-581 (581). Data shown is n=4 and were validated in 2 separated experiments. Significance testing was performed by Student *t* test at day 4 and no significant difference was observed in each conditions (*P*=0.0533 between DMSO and 250nM group). C, Invadopodia formation analysis in C8161.9 cells with control DMSO and TAO3 inhibitor, SBI-029 (029). D, Mouse weight at day 21 after treatment of DMSO or SBI-581 (581). P>0.05 unless other specified; *, *P*<0.05.

**Supplementary Figure S4.**
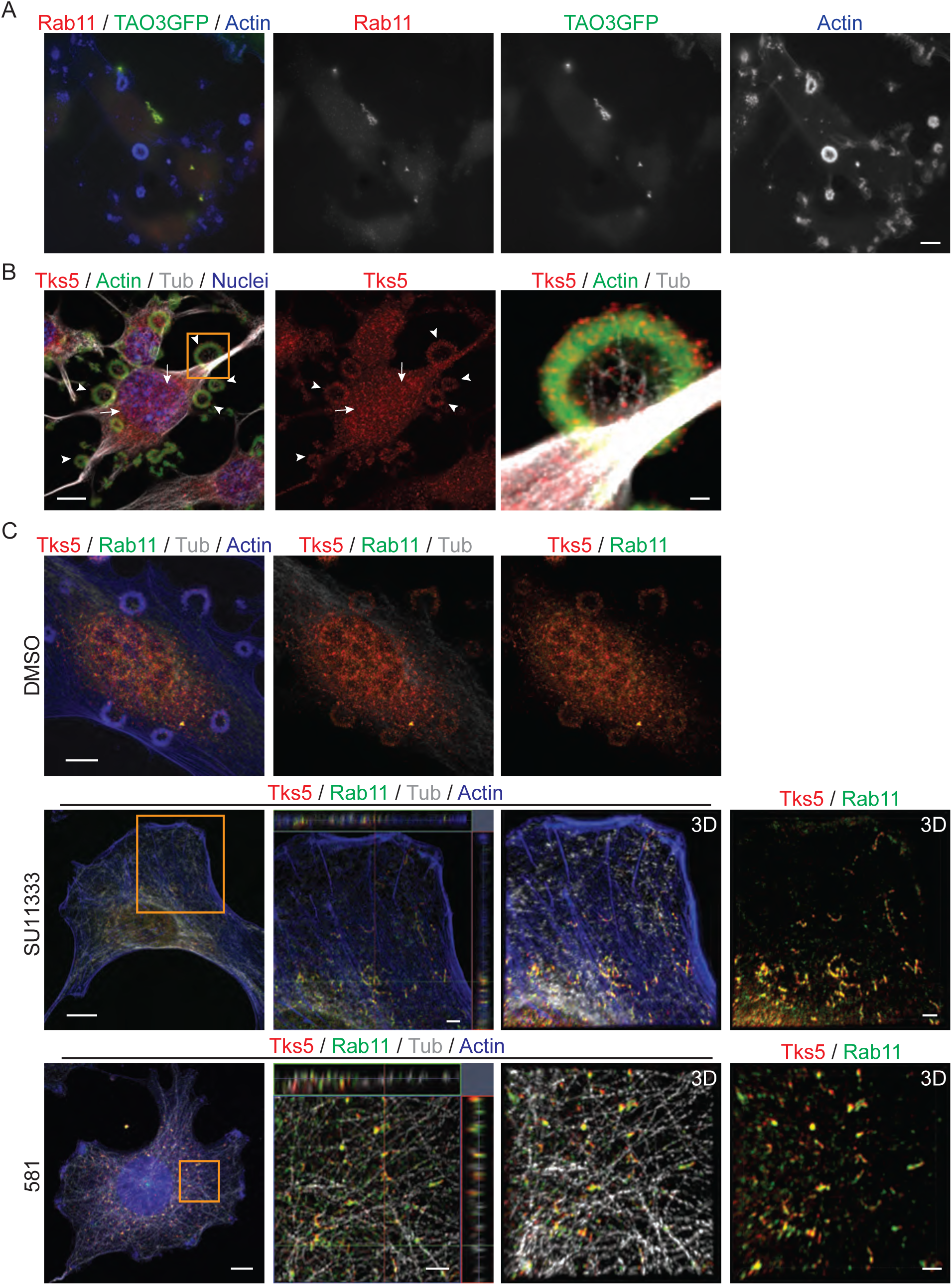
Localization of TAO3 and Tks5 at Rab11+ endosomal vesicles. A, TAO3-GFP (green) was overexpressed in Src3T3 cells and stained by Rab11 (red) and Actin (phalloidin, blue). B, Distribution of Tks5. Staining of Tks5 (red), actin (green), tubulin (gray) and nuclei (blue) in Src3T3 cells. Images were processed by maximum intensity projection. Arrowheads indicate invadopodial positioning of Tks5. Arrows indicate endosomal positioning of Tks5. Magnified area (orange square in left) was shown (right). C, Accumulation of Tks5 at Rab11+ endosomal vesicles. Staining of Tks5 (red), Rab11 (green), tubulin (gray) and actin (phalloidin, blue) in Src3T3 cells. Images were processed by maximum intensity projection (top three panels) with different combination of channels indicated in each image. Invadopodia formation was inhibited by SU11333 (middle three) or SBI-581 (bottom three) and stained. Magnified images (orange square in left) were shown as orthogonal view (second left) or 3D reconstruction by Imaris software (3D, right two with different combination of channels shown in each images). Scale bars, 10 μm (A, B left and C left), 1 μm (B right) and 2 μm (C second left and right).

**Supplementary Figure S5.**
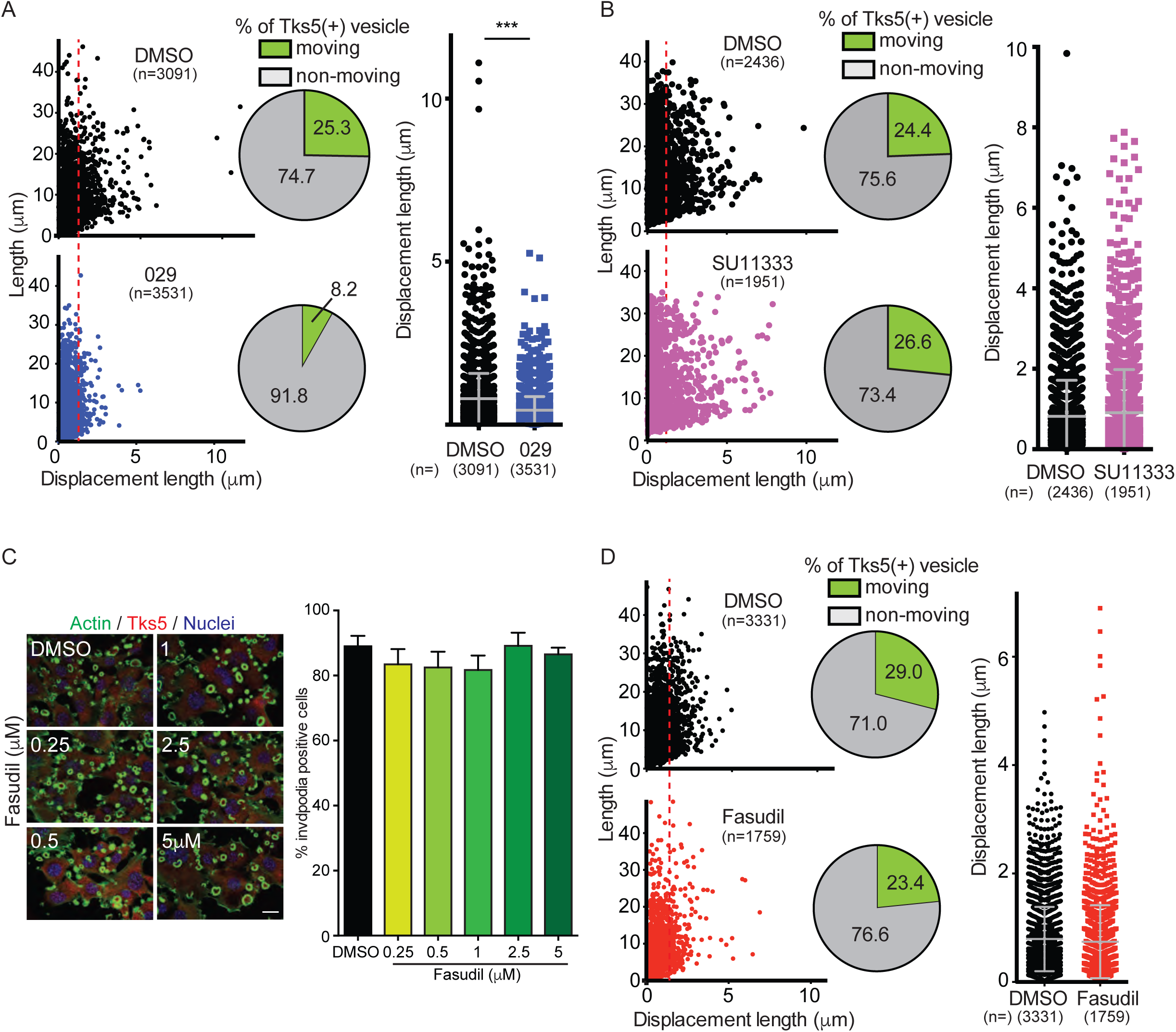
Trafficking of Tks5+ vesicles (A, B, D) was captured by time-lapse imaging (200ms/image for 1 min, total 300 images/film) in Src3T3 cells expressing Tks5-mCherry and YFP-tubulin. Src3T3 cells expressing Tks5-mCherry and YFP-tubulin were treated by DMSO, SBI-029 (A, 029, 100nM), SU11333 (B, 1μM) or Fasudil (D, 1μM) and moving of Tks5 signal was analyzed by plotting graph of length/displacement length (left) with percentage of Tks5+ vesicle moving (pie chart) and displacement length (right). Data shown are n=3 (unless other specified in figure) and were validated in 2 or more separated experiments. *P*>0.05 unless other specified; ***, *P*<0.001. C, Invadopodia formation analysis in Src3T3 cells with control DMSO and ROCK2 inhibitor, Fasudil. Representative images (left) and percentage of invadopodia positive cells (right). Immunofluorescence staining of invadopodia by actin (phalloidin, green) and TKS5 (red), and Hoechst to denote nuclei of cells. Data shown are n=5 in each experimental group and were validated in 2 or more separate experiments. Scale bar, 20μm.

**Supplementary Figure S6.**
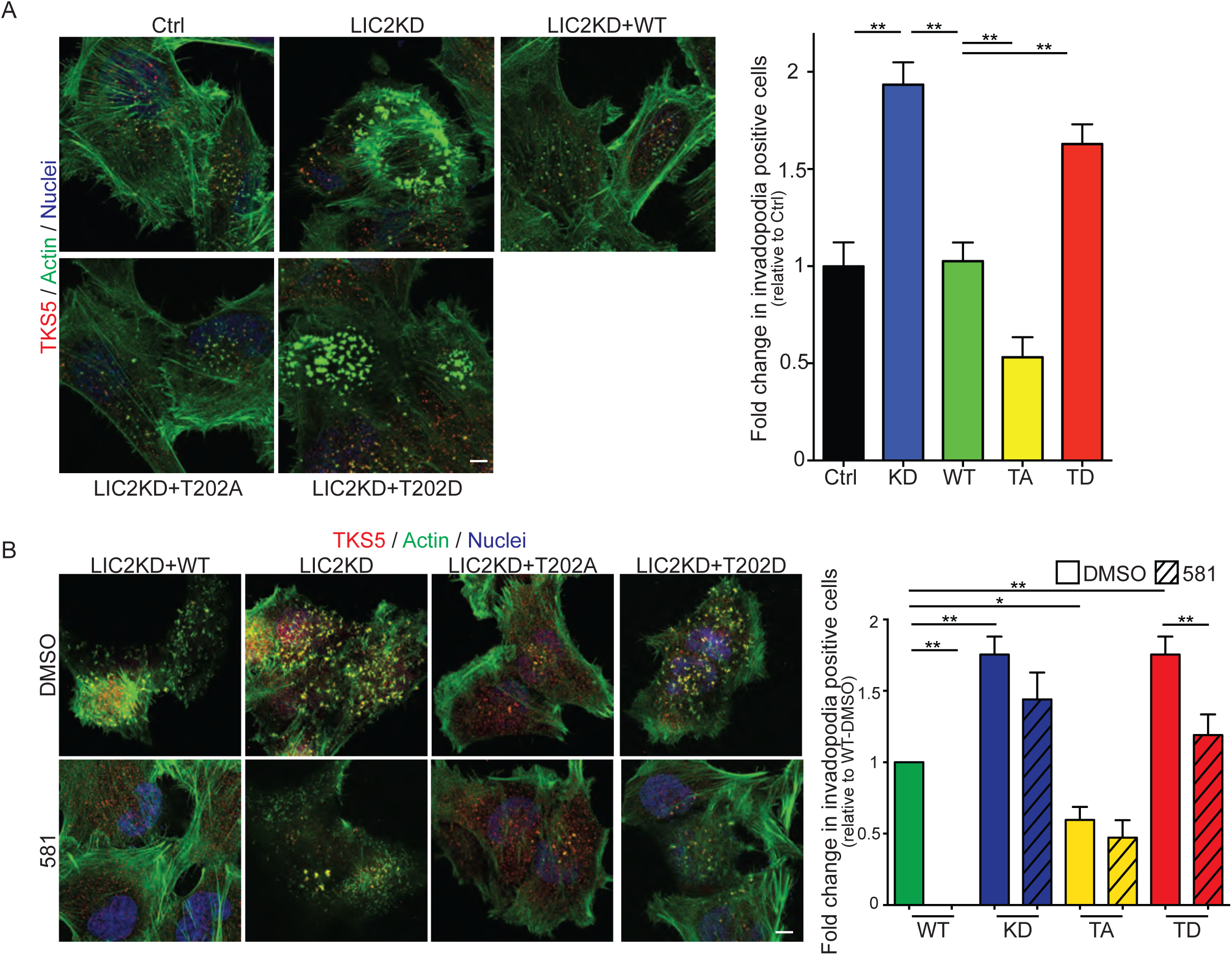
LIC2 regulates invadopodia formation in melanoma cell line, WM793 (A, B). Invadopodia formation (left: representative images, right: percentage or relative invadopodia positive cells) in WM793 cells with control (Ctrl), LIC2 knockdown (KD), rescued expression of shRNA-resistant-LIC2 wild-type (WT) or threonine 202 point mutations (T202A;TA or T202D;TD). Immunofluorescence staining of invadopodia by actin (phalloidin, green) and Hoechst to denote nuclei of cells. B, The cells are cultured on high-density collagen with or without TAO3 inhibitor, SBI-581 (581). Data shown are n=5 in each experimental group and were validated in 2 or more separated experiments. *P*>0.05 unless other specified; **, *P*<0.01; ***, *P*<0.001.

**Supplementary Table S1.** Outlier analysis for TAOK3 based on Gene Rank Threshold of 10%. The sample percentile indicates the subset of samples with over-expressed levels of gene expression (e.g., 75th percentile considers the top 25 percent of the samples for expression of TAOK3). The Gene Rank is % Rank based on over-expression of TAOK3 in that study. A high gene rank (which is based on the COPA score below) indicates there are few genes that are more significant within the dataset (i.e., have a higher COPA score). COPA is the Cancer Outlier Profile Analysis score, the transformed expression value for the outlier analysis. A higher COPA score indicates a more significant outlier profile.

| Tumor type | Study | Sample Percentile | Gene Rank% | COPA |
| --- | --- | --- | --- | --- |
| Bladder | Lee | 95<sup>th</sup> | 1% | 33.17 |
| Breast | Finak | 95<sup>th</sup> | 2% | 17.33 |
| Breast | Farmer | 75<sup>th</sup> | 3% | 1.93 |
| Breast (2) | Schuetz | 75<sup>th</sup> | 3% | 2.91 |
| Breast | Loi | 95<sup>th</sup> | 9% | 4.76 |
| Melanoma | Harlin | 95<sup>th</sup> | 7% | 5.60 |
| Melanoma | Bogunivic | 75<sup>th</sup> | 3% | 1.91 |
| Head | Ginos | 75<sup>th</sup> | 8% | 1.50 |
| Head | Slebos | 95<sup>th</sup> | 6% | 6.71 |
| Glioblastoma | Dong | 95<sup>th</sup> | 5% | 6.83 |
| Colon | Lin | 90<sup>th</sup> | 7% | 3.05 |
|  |  | 95<sup>th</sup> | 9% | 4.17 |
| Esophageal | Shimokuni (Cell Line 2) | 75<sup>th</sup> | 2% | 2.33 |
| Prostate | LaTuillipe | 75<sup>th</sup> | 4% | 1.81 |
| Leukemia | SaiyaCork | 75<sup>th</sup> | 1% | 3.55 |
|  |  | 90<sup>th</sup> | 6% | 5.77 |
| Leukemia | Maia | 95<sup>th</sup> | 5% | 4.43 |
| Leukemia | Cario | 95<sup>th</sup> | 5% | 5.51 |
| Leukemia | Falt | 95<sup>th</sup> | 10% | 3.74 |
| Lymphoma | Braggio | 95<sup>th</sup> | 3% | 17.98 |
| Lymphoma 3 | Rosenwald | 95<sup>th</sup> | 7% | 4.12 |
| Renal | Gumz | 75<sup>th</sup> | 4% | 2.08 |
| Liver | Archer | 95<sup>th</sup> | 3% | 5.05 |
| Liver | Mas | 95<sup>th</sup> | 4% | 5.14 |
| Lung | Tse (Cell Line) | 75<sup>th</sup> | 5% | 1.82 |
| Mesothelioma | LopezRios | 95<sup>th</sup> | 7% | 5.41 |
| Ovarian | Lu | 75<sup>th</sup> | 2% | 2.07 |
| Ovarian | Hendrix | 75<sup>th</sup> | 4% | 1.50 |
| Pancreas 2 | IacobuzioDonahue | 90<sup>th</sup> | 8% | 3.28 |
| Sarcoma | Ohali | 75<sup>th</sup> | 2% | 3.99 |
|  |  | 90<sup>th</sup> | 8% | 4.64 |

**Supplementary Table S2.**
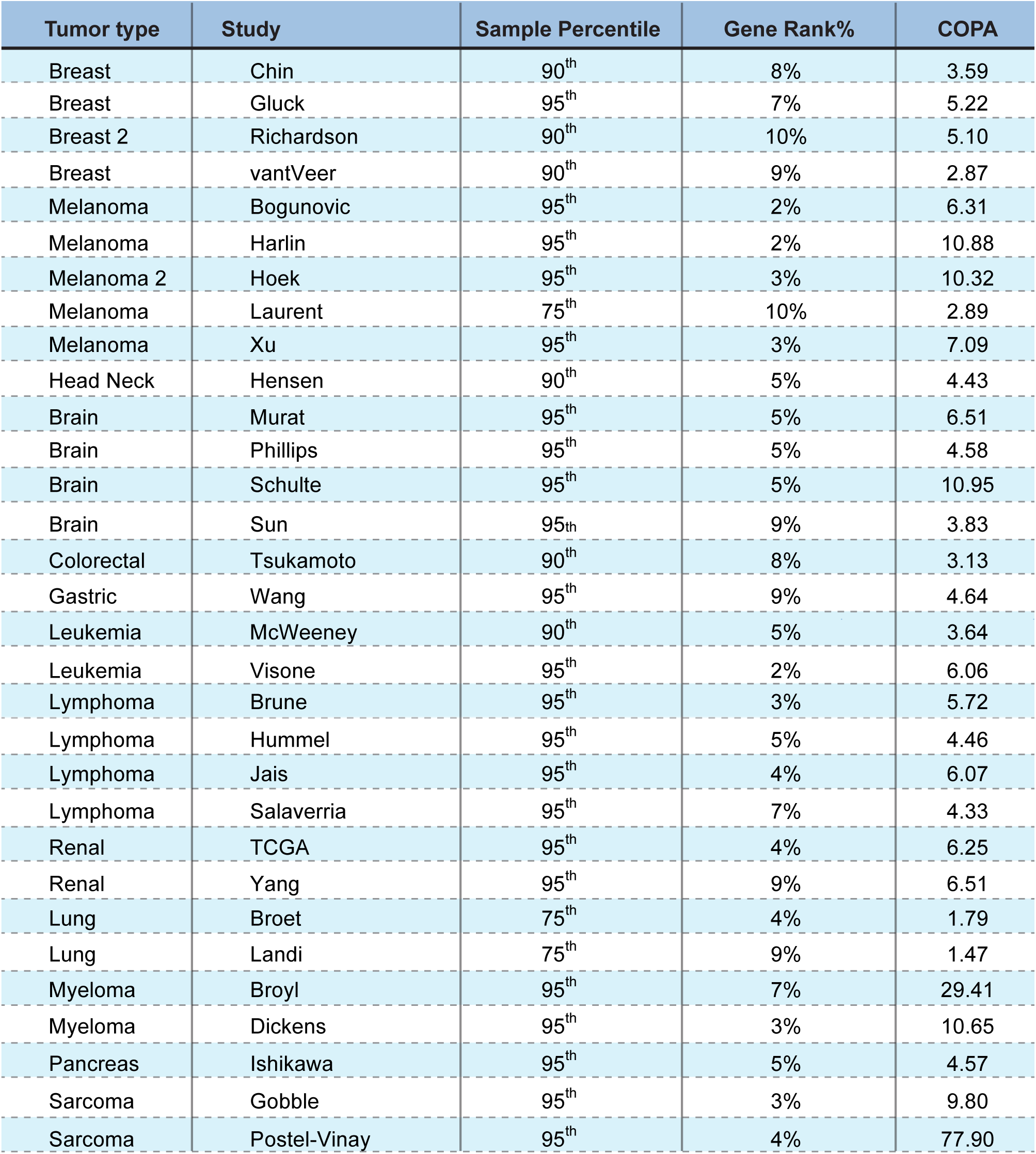
Outlier analysis for PAK6 based on Gene Rank Threshold of 10%. The sample percentile indicates the subset of samples with over-expressed levels of gene expression (e.g., 75th percentile considers the top 25 percent of the samples for expression of PAK6). The Gene Rank is % Rank based on over-expression of PAK6 in that study. A high gene rank (which is based on the COPA score below) indicates there are few genes that are more significant within the dataset (i.e., have a higher COPA score). COPA is the Cancer Outlier Profile Analysis score, the transformed expression value for the outlier analysis. A higher COPA score indicates a more significant outlier profile.

| Tumor type | Study | Sample Percentile | Gene Rank% | COPA |
| --- | --- | --- | --- | --- |
| Breast | Chin | 90<sup>th</sup> | 8% | 3.59 |
| Breast | Gluck | 95<sup>th</sup> | 7% | 5.22 |
| Breast 2 | Richardson | 90<sup>th</sup> | 10% | 5.10 |
| Breast | vantVeer | 90<sup>th</sup> | 9% | 2.87 |
| Melanoma | Bogunovic | 95<sup>th</sup> | 2% | 6.31 |
| Melanoma | Harlin | 95<sup>th</sup> | 2% | 10.88 |
| Melanoma 2 | Hoek | 95<sup>th</sup> | 3% | 10.32 |
| Melanoma | Laurent | 75<sup>th</sup> | 10% | 2.89 |
| Melanoma | Xu | 95<sup>th</sup> | 3% | 7.09 |
| Head Neck | Hensen | 90<sup>th</sup> | 5% | 4.43 |
| Brain | Murat | 95<sup>th</sup> | 5% | 6.51 |
| Brain | Phillips | 95<sup>th</sup> | 5% | 4.58 |
| Brain | Schulte | 95<sup>th</sup> | 5% | 10.95 |
| Brain | Sun | 95<sup>th</sup> | 9% | 3.83 |
| Colorectal | Tsukamoto | 90<sup>th</sup> | 8% | 3.13 |
| Gastric | Wang | 95<sup>th</sup> | 9% | 4.64 |
| Leukemia | McWeeney | 90<sup>th</sup> | 5% | 3.64 |
| Leukemia | Visone | 95<sup>th</sup> | 2% | 6.06 |
| Lymphoma | Brune | 95<sup>th</sup> | 3% | 5.72 |
| Lymphoma | Hummel | 95<sup>th</sup> | 5% | 4.46 |
| Lymphoma | Jais | 95<sup>th</sup> | 4% | 6.07 |
| Lymphoma | Salaverria | 95<sup>th</sup> | 7% | 4.33 |
| Renal | TCGA | 95<sup>th</sup> | 4% | 6.25 |
| Renal | Yang | 95<sup>th</sup> | 9% | 6.51 |
| Lung | Broet | 75<sup>th</sup> | 4% | 1.79 |
| Lung | Landi | 75<sup>th</sup> | 9% | 1.47 |
| Myeloma | Broyl | 95<sup>th</sup> | 7% | 29.41 |
| Myeloma | Dickens | 95<sup>th</sup> | 3% | 10.65 |
| Pancreas | Ishikawa | 95<sup>th</sup> | 5% | 4.57 |
| Sarcoma | Gobble | 95<sup>th</sup> | 3% | 9.80 |
| Sarcoma | Postel-Vinay | 95<sup>th</sup> | 4% | 77.90 |

**Supplementary Table S3.** Outlier analysis for CAMK1 based on Gene Rank Threshold of 10%. The sample percentile indicates the subset of samples with over-expressed levels of gene expression (e.g., 75th percentile considers the top 25 percent of the samples for expression of CAMK1). The Gene Rank is % Rank based on over-expression of CAMK1 in that study. A high gene rank (which is based on the COPA score below) indicates there are few genes that are more significant within the dataset (i.e., have a higher COPA score). COPA is the Cancer Outlier Profile Analysis score, the transformed expression value for the outlier analysis. A higher COPA score indicates a more significant outlier profile.

| Tumor type | Study | Sample Percentile | Gene Rank% | COPA |
| --- | --- | --- | --- | --- |
| Bladder 2 | Blaveri | 90<sup>th</sup> | 6% | 2.96 |
| Breast | Kreike | 75<sup>th</sup> | 7% | 1.48 |
| Breast 2 | Perou | 95<sup>th</sup> | 8% | 4.63 |
| Melanoma | Laurent | 95<sup>th</sup> | 9% | 26.30 |
| Head-Neck | Peng | 75<sup>th</sup> | 5% | 1.49 |
| Brain | Freije | 90<sup>th</sup> | 10% | 2.98 |
| Brain | Sun | 95<sup>th</sup> | 8% | 3.87 |
| Brain | Janoueix- | 90<sup>th</sup> | 8% | 33.00 |
| Colorectal 3 | Jorissen | 95<sup>th</sup> | 6% | 4.63 |
| Esophagus | Kim | 75<sup>th</sup> | 7% | 1.65 |
| Prostate | Varambally | 95<sup>th</sup> | 8% | 7.10 |
| Prostate | Singh | 95<sup>th</sup> | 1% | 18.92 |
| Cervix | Biewenga | 75<sup>th</sup> | 8% | 1.62 |
| Leukemia | Andersson | 95<sup>th</sup> | 5% | 4.44 |
| Leukemia | Armstrong | 95<sup>th</sup> | 5% | 15.91 |
| Leukemia | Balgobind | 95<sup>th</sup> | 4% | 5.56 |
| Leukemia | Haslinger | 95<sup>th</sup> | 7% | 4.10 |
| Leukemia 2 | Metzeler | 95<sup>th</sup> | 1% | 9.37 |
|  |  | 95<sup>th</sup> | 4% | 4.89 |
| Leukemia | McWeeney | 75<sup>th</sup> | 5% | 1.73 |
| Leukemia | TCGA | 95<sup>th</sup> | 9% | 3.52 |
| Leukemia | Tsutsumi | 95<sup>th</sup> | 1% | 72.61 |
| Leukemia | Valk | 95<sup>th</sup> | 10% | 3.73 |
| Leukemia | Visone | 95<sup>th</sup> | 4% | 4.79 |
| Leukemia | Wouters | 90<sup>th</sup> | 8% | 3.41 |
| Lymphoma | Chng | 90<sup>th</sup> | 8% | 3.39 |
| Lymphoma | Compagno | 75<sup>th</sup> | 8% | 1.45 |
| Lymphoma | Dave | 95<sup>th</sup> | 6% | 4.22 |
| Lymphoma | Hummel | 95<sup>th</sup> | 1% | 7.29 |
| Lymphoma | Klapper | 95<sup>th</sup> | 3% | 7.49 |
| Lymphoma | Lenz | 95<sup>th</sup> | 10% | 3.50 |
| Lymphoma | Lossos | 95<sup>th</sup> | 10% | 5.04 |
| Lymphoma | Piccaluga | 95<sup>th</sup> | 2% | 10.58 |
| Lymphoma | Rosenwald | 90<sup>th</sup> | 4% | 3.09 |
|  |  | 95<sup>th</sup> | 7% | 3.47 |
| Lymphoma | Salaverria | 95<sup>th</sup> | 3% | 5.20 |
| Lymphoma | Shaknovich | 95<sup>th</sup> | 8% | 3.85 |
| Renal | Cutcliffe | 95<sup>th</sup> | 4% | 5.37 |
| Renal | Jones | 75<sup>th</sup> | 7% | 1.64 |
| Renal | Vasselli | 90<sup>th</sup> | 8% | 3.19 |
| Liver | Wurmbach | 75<sup>th</sup> | 7% | 2.68 |
| Liver | Ye | 95<sup>th</sup> | 4% | 4.83 |
| Ovarian | Lu | 95<sup>th</sup> | 9% | 5.56 |
| Ovarian | Hendrix | 95<sup>th</sup> | 6% | 4.33 |
| Pancreas 2 | Iacobuzio-Donahue | 95<sup>th</sup> | 6% | 4.95 |
| Sarcoma | Chibon | 75<sup>th</sup> | 7% | 5.503 |

**Supplementary Table S4.**
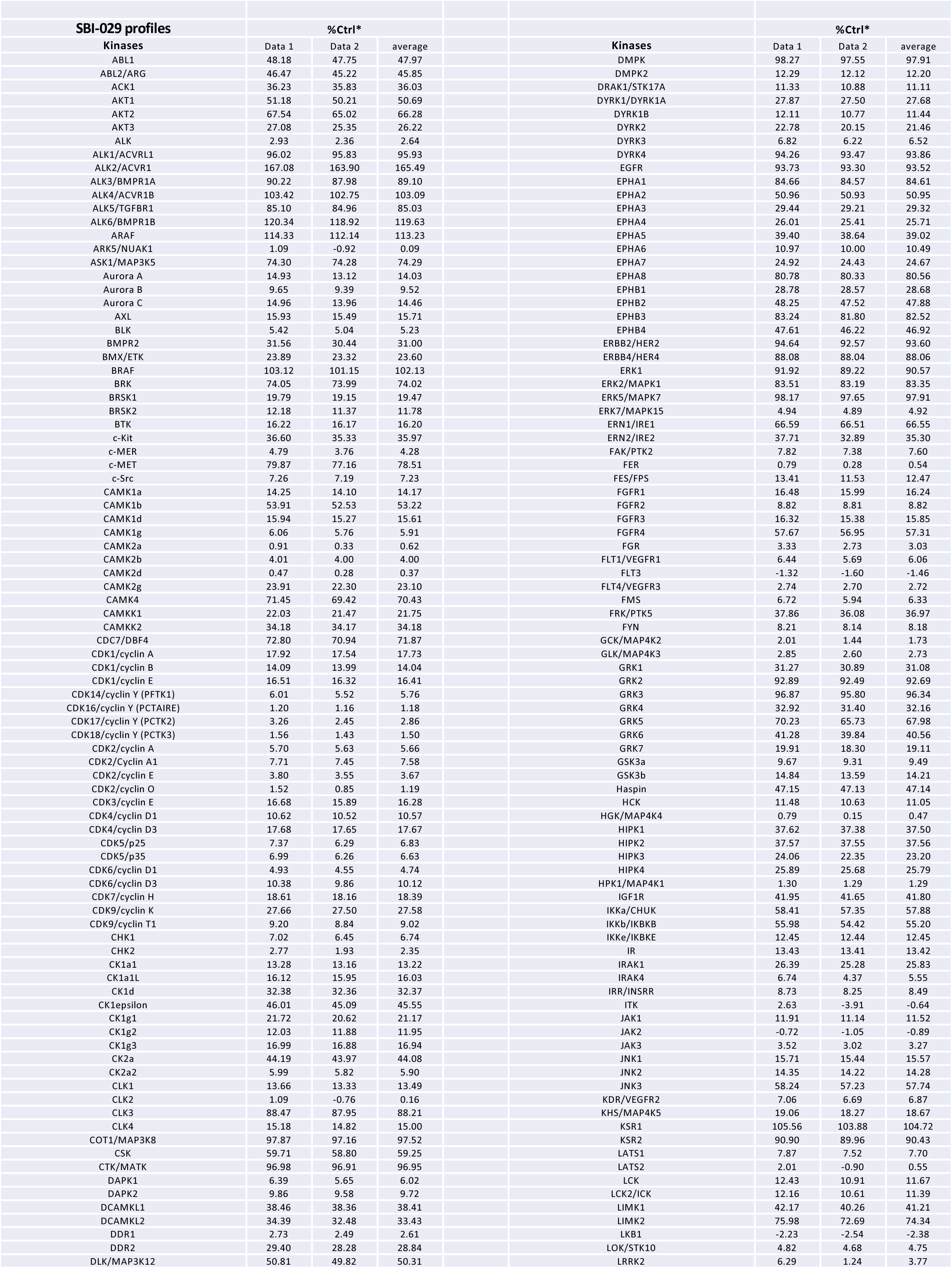
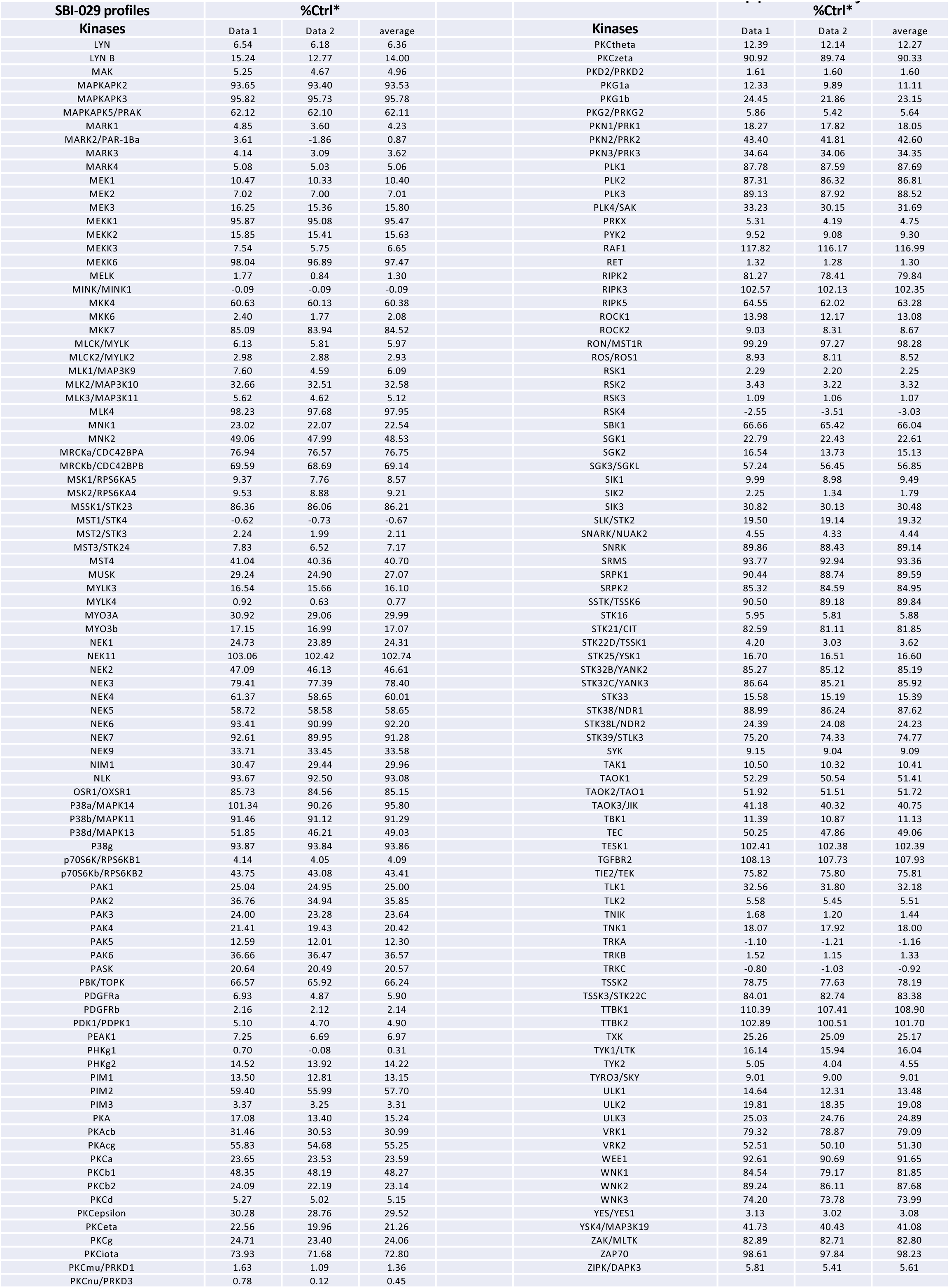
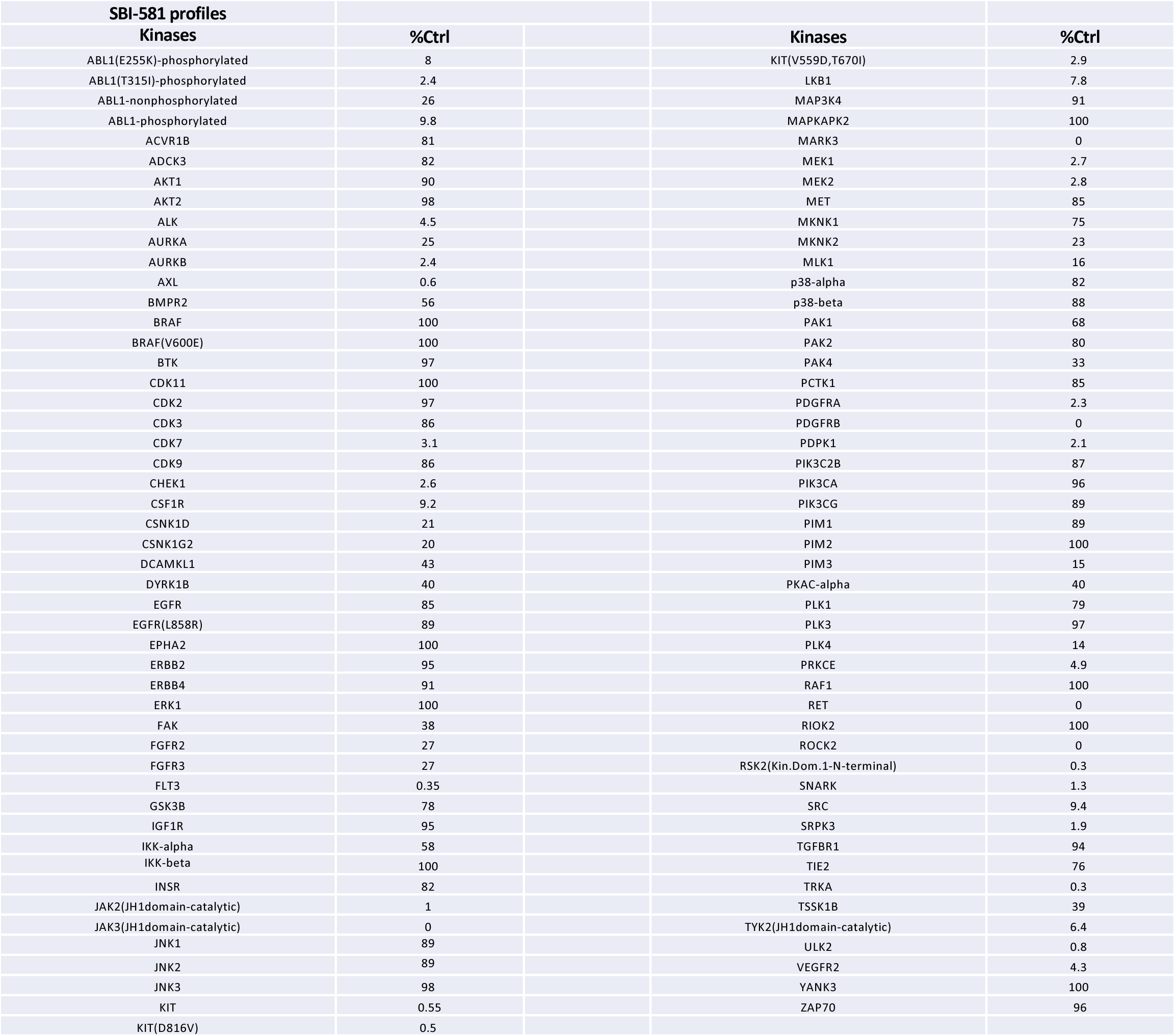
**SBI compound profiles.** List of kinase activities with TAO3 inhibitors, SBI-029 (page1 and 2) or SBI-581 (page3) *%Ctrl compared to kinase dead form.

**Supplementary Table S5.**
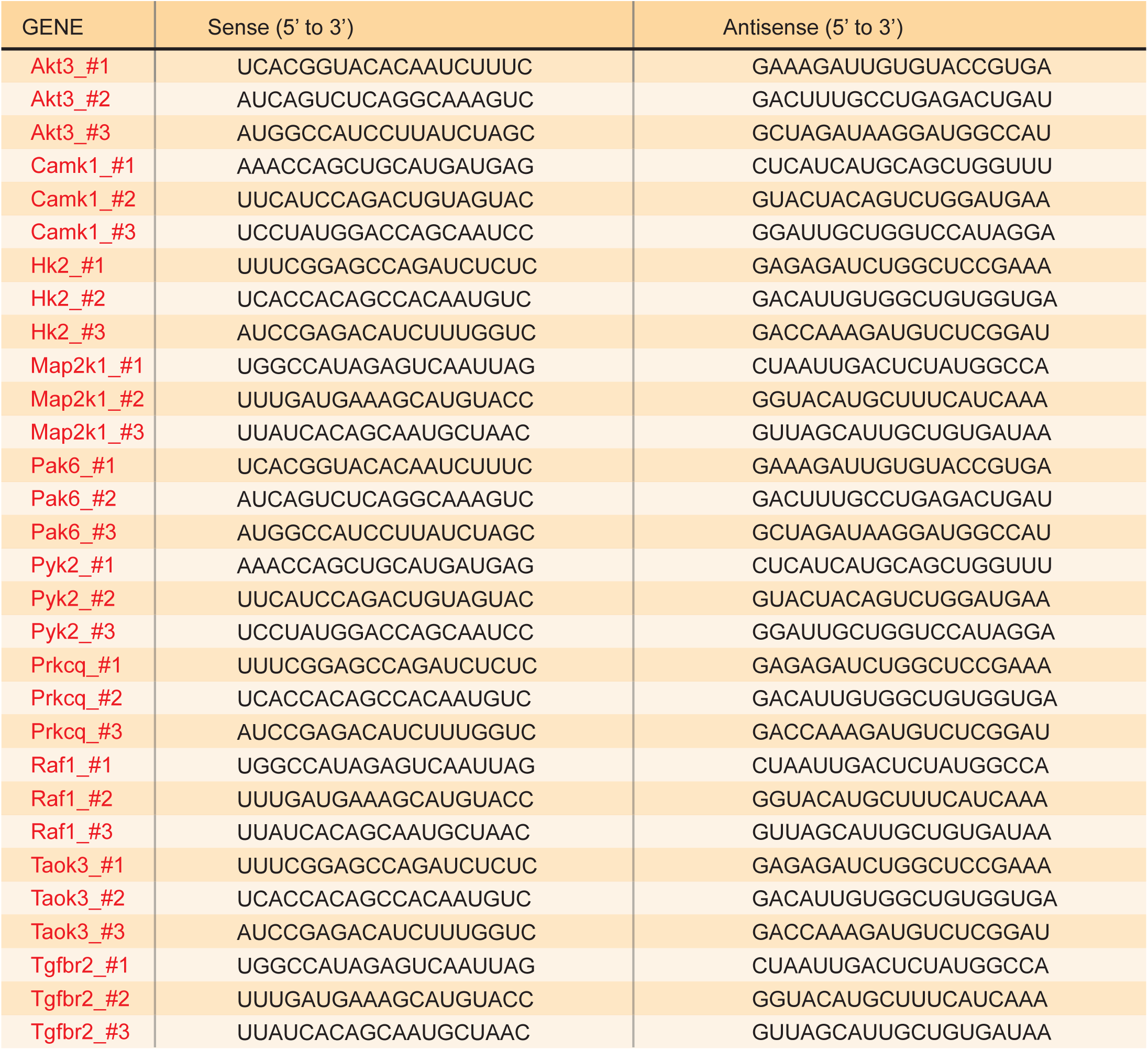
List of siRNAs for validation assay (related to Figure 1, Table 1)

